# Rapid and strain-specific resistance evolution of *Staphylococcus aureus* against inhibitory molecules secreted by *Pseudomonas aeruginosa*

**DOI:** 10.1101/2021.08.10.455797

**Authors:** Selina Niggli, Lukas Schwyter, Lucy Poveda, Jonas Grossmann, Rolf Kümmerli

**Affiliations:** Department of Quantitative Biomedicine, University of Zurich, Winterthurerstrasse 190, 8057 Zurich, Switzerland; Functional Genomics Center Zurich, ETH Zurich and University of Zurich, Winterthurerstrasse 190, 8057 Zurich, Switzerland

## Abstract

*Pseudomonas aeruginosa* and *Staphylococcus aureus* frequently occur together in polymicrobial infections, and there is evidence that their interactions negatively affect disease outcome in patients. At the molecular level, interactions between the two bacterial species are well-described, with *P. aeruginosa* usually being the dominant species suppressing *S. aureus* through a variety of inhibitory molecules. However, in chronic infections the two species interact over prolonged periods of time, and *S. aureus* might be able to evolve resistance against inhibitory molecules deployed by *P. aeruginosa*. Here, we used experimental evolution to test this hypothesis by exposing three different *S. aureus* strains (Cowan I, 6850, JE2) to the growth-inhibitory supernatant of *P. aeruginosa* PAO1 over 30 days. Prior to evolution, we found that *S. aureus* strains were inhibited by secreted compounds regulatorily controlled by the PQS quorum-sensing system of *P. aeruginosa*. Following evolution, *S. aureus* strains were no longer inhibited: we observed that phenotypic adaptations were strain-specific and involved the up-regulation of virulence traits, such as staphyloxanthin production and the formation of small colony variants. At the genetic level, mutations in membrane transporters were the most frequent evolutionary targets. Our work indicates that adaptations of *S. aureus* to co-infecting pathogens occurs rapidly and involves both virulence traits and membrane transporters involved in drug resistance. Thus, pathogen evolution could promote species co-existence, complicate treatment options and therefore worsen disease outcome.

## Introduction

Bacterial infections are frequently caused by multiple species (1–3) and such polymicrobial infections can be more severe than the respective mono-species infections (4–7). Because of the detrimental effects for patients, there is high interest in understanding the mechanisms of bacterial interactions and how they affect ecological dynamics between co-infecting species. Interactions between the two opportunistic human pathogens *Pseudomonas aeruginosa* (PA) and *Staphylococcus aureus* (SA) have attracted particularly high attention (8, 9). This is because they are major pathogens and often co-occur in the lungs of cystic fibrosis (CF) patients, and in burn and chronic wound infections (10–13). Co-infections with PA and SA can be associated with more severe disease outcomes in humans (13, 14), and studies in animal models confirmed increased virulence and compromised antibiotic treatment options during co-infections (15–17).

Laboratory studies established a solid mechanistic understanding of how the two species interact, and generally agree that the relationship between PA and SA is antagonistic (18–20). Overall, PA seems to dominate the interactions through the production and secretion of a variety of inhibitory molecules, such as the staphylolytic protease LasA, siderophores, and the respiratory chain inhibitors HQNO and phenazines (21–26). There is also increasing knowledge on ecological factors that influence interactions patterns. For example, during early childhood, CF lungs are first colonized by SA followed by PA later on, with SA abundance tending to decrease when PA increases (27). In the context of wound infections, we also have an increased understanding about the spatial localization of the two species, where it seems that PA and SA occupy different niches (28, 29).

In contrast to these mechanistic and ecological insights, we know little about whether interactions between PA and SA can evolve (30). Species co-evolution is likely to occur in chronic infections, where pathogens repeatedly interact, and is predicted to alter disease parameters and treatment strategies. In the context of interactions between PA and SA, we hypothesize that SA could adapt and become resistant to PA inhibitory molecules. Such adaptation could in turn foster co-existence of the two species, potentially contribute to the clinically observed higher virulence, increased morbidity and treatment complications (13, 14).

Here, we use a combination of experimental evolution, phenotypic screening, and whole-genome sequencing of evolved clones to test our hypothesis on resistance evolution. Specifically, we first conducted a series of experiments with three different clinical SA strains to show that all of them are inhibited by a secreted compound present in the supernatant and regulated by the PQS quorum-sensing system of the PA strain PAO1. We then exposed the three SA strains to either the PA supernatant (containing the inhibitory molecules, mixed with fresh medium) or a control medium for 30 consecutive days, by transferring evolving cultures every 48 hours to fresh conditions. Following experimental evolution, we screened evolved SA clones from replicated populations for resistance phenotypes and explored whether virulence traits were under selection. Finally, we sequenced the whole genome of 150 clones to determine the genetic basis of SA resistance evolution against inhibitory molecules from PA.

## Results

### Growth of SA strains is reduced in the presence of PA supernatant

Previous work showed that PA can inhibit SA via a diverse set of secreted molecules, including proteases, biosurfactants, siderophores, and toxic compounds (8). To test whether our three SA strains (Cowan I, 6850 and JE2, Table S1) are also inhibited by PA, we exposed them to the sterile-filtered supernatant of PA PAO1, harvested from overnight cultures (30% supernatant in 70% fresh tryptic soy broth, TSB). We found that all three SA strains were negatively affected by the PA supernatant compared to growth under control condition (30% NaCl in 70% TSB, Figure 1). Exposure to PA supernatant significantly reduced overall growth performance (Figure S1a, ANOVA on growth integrals followed by Tukey’s HSD pairwise comparisons: p_adj_ < 0.0001 for all SA strains), and specifically extended the lag phase of all SA strains (Figure S1b). We also observed SA strain-specific responses (Figure 1). Cowan I featured an intermediate extension of the lag phase (10.3 hours compared to 6.6 hours in the control medium), a premature stationary phase, followed by a decrease in OD_600_ possibly indicating cell death. 6850 suffered from an extremely long lag phase extension (19.7 hours vs. 6.9 hours), also showed a premature growth stop, but no OD_600_ decline. JE2 was least affected by the PA supernatant. Its lag phase extension was not as pronounced (8.4 hours vs. 5.6 hours), and it grew steadily afterwards, albeit to a lower yield than the control.

**Figure 1.**
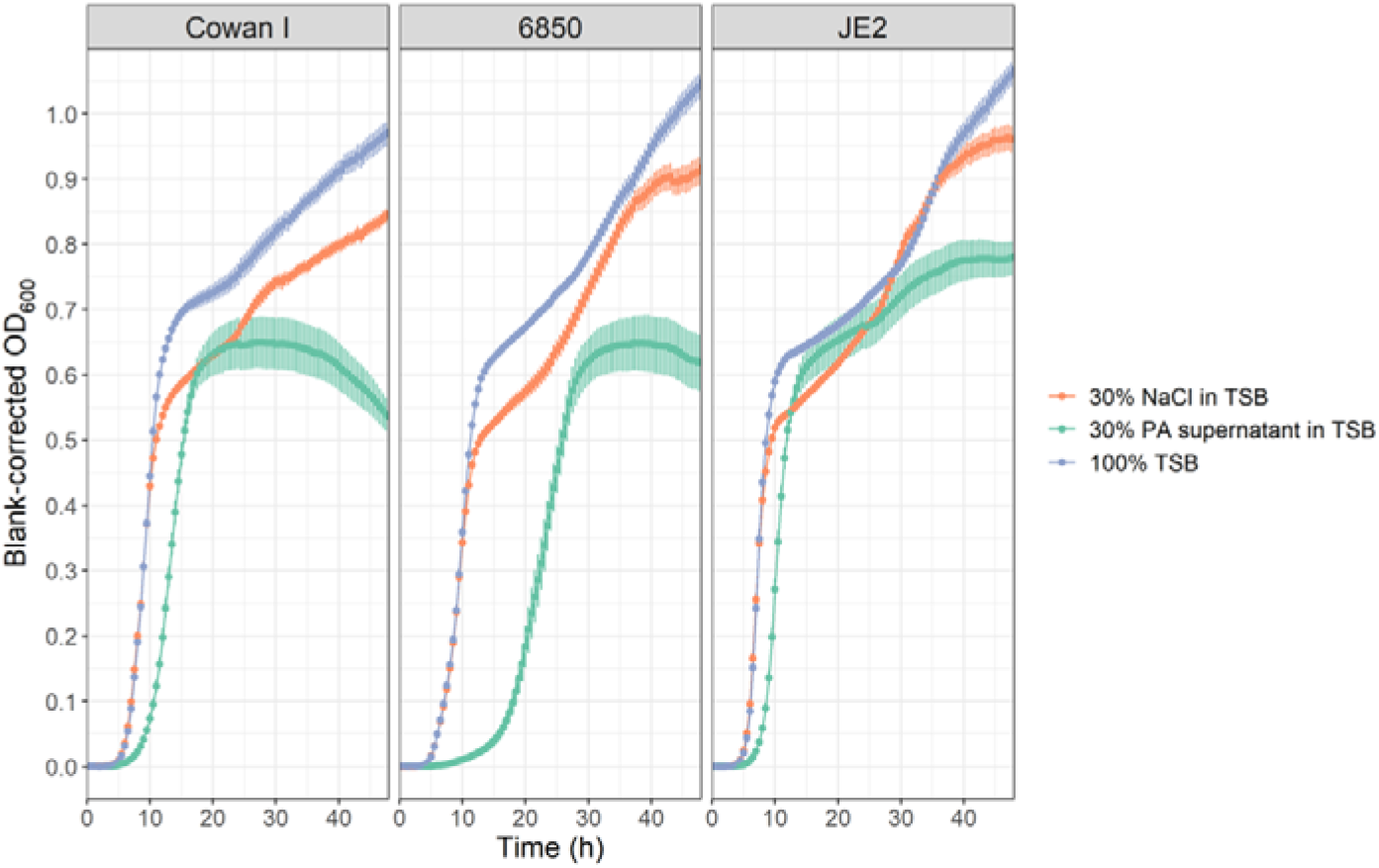
Supernatant of PA inhibits growth of three different SA strains (Cowan I, 6850, JE2). Growth trajectories of SA strains were followed over 48 hours at 37 °C under three different conditions: 30% NaCl in TSB (control 1), 30% PA supernatant in TSB (experimental treatment) and 100% TSB (control 2). Curves show the average growth (± standard error) across seven independent experiments each featuring four replicates per strain and condition. See Figure S1 for a statistical comparison of growth parameters extracted from these curves. The results from all statistical analyses can be found in the Statistics Source File.

We also conducted the reciprocal experiments by exposing PA to the supernatants of the three SA strains but found no inhibition (Figure S2). Taken together, our supernatant assays reveal unilateral inhibition: PA produces inhibitory molecules that negatively affect the growth of genetically different SA strains.

### The PQS pathway is involved in compromising SA growth

To identify possible PA pathways or traits involved in the inhibition of SA, we screened a small panel of seven PA candidate mutants and checked whether the inhibitory effects of their supernatants towards SA were altered. We found one hit: the supernatant of PAO1Δ*pqsA* did no longer inhibit SA (Table 1, Figure S3).

**Table 1.** Relative effects of supernatants from wildtype and mutant *P. aeruginosa* (PA) strains on *S. aureus* (SA) growth.

| PA supernatant donor strain | Phenotype of mutant strain | Relative growth* of Cowan I in PA supernatant | Relative growth* of 6850 in PA supernatant | Relative growth* of JE2 in PA supernatant |
| --- | --- | --- | --- | --- |
| PAO1 wild type | wild type | 0.78 (± 0.20) | 0.60 (± 0.23) | 0.87 (± 0.18) |
| MPAO1 wild type | wild type | 0.75 (± 0.23) | 0.59 (± 0.21) | 0.72 (± 0.18) |
| PAO1ΔlasR | Las quorum sensing signal blind | 0.34 (± 0.08) | 0.38 (± 0.10) | 0.50 (± 0.19) |
| PAO1ΔrhII | Rhl quorum sensing signal negative | 0.65 (± 0.05) | 0.45 (± 0.11) | 0.67 (± 0.05) |
| PAO1ΔrhIR | Rhl quorum sensing signal blind | 0.70 (± 0.13) | 0.60 (± 0.05) | 0.70 (± 0.03) |
| PAO1ΔrhIA | Rhamnolipid null mutant | 0.61 (± 0.11) | 0.54 (± 0.15) | 0.68 (± 0.07) |
| MPAO1ΔpqsA | Alkylquinolone null mutant | **1.22 (± 0.16)** | **1.22 (± 0.10)** | **1.05 (± 0.12)** |
| PAO1ΔpvdDΔpchEF | Siderophore null mutant | 0.63 (± 0.01) | 0.23 (± 0.04) | 0.68 (± 0.02) |
| PAO1ΔlecA | Lectin null mutant | 0.67 (± 0.02) | 0.42 (± 0.01) | 0.83 (± 0.06) |

\*The relative supernatant effect ( ± standard deviation) is expressed as the growth integral of SA over 48 hours in 30% PA supernatant + TSB divided by the growth integral of SA in 30% NaCl + TSB. Values < 1 indicate growth inhibition, while values > 1 (in bold) indicate cases with no inhibition.

PqsA is an anthranilate-coenzyme A ligase that plays a role in PA cell-to-cell communication (quorum sensing) (31). It is directly involved in the synthesis of a family of secondary metabolites, including 4-hydroxy-2-heptylquinoline (HHQ), 2-n-heptyl-4-hydroxyquinoline N-oxide (HQNO) and the *Pseudomonas* quinolone signal (PQS). HHQ, HQNO and PQS have previously been suggested to inhibit a variety of bacterial and fungal species (24, 32–34). Here, we tested whether they also inhibit our SA strains by focusing on PQS as one of the molecules synthetized by PqsA. We exposed all three SA strains to increasing concentrations of synthetic PQS and found a dose-dependent reduction in growth (Figure 2). This indicates that PQS is involved in the SA growth inhibition that we observed in our supernatant assays.

**Figure 2.**
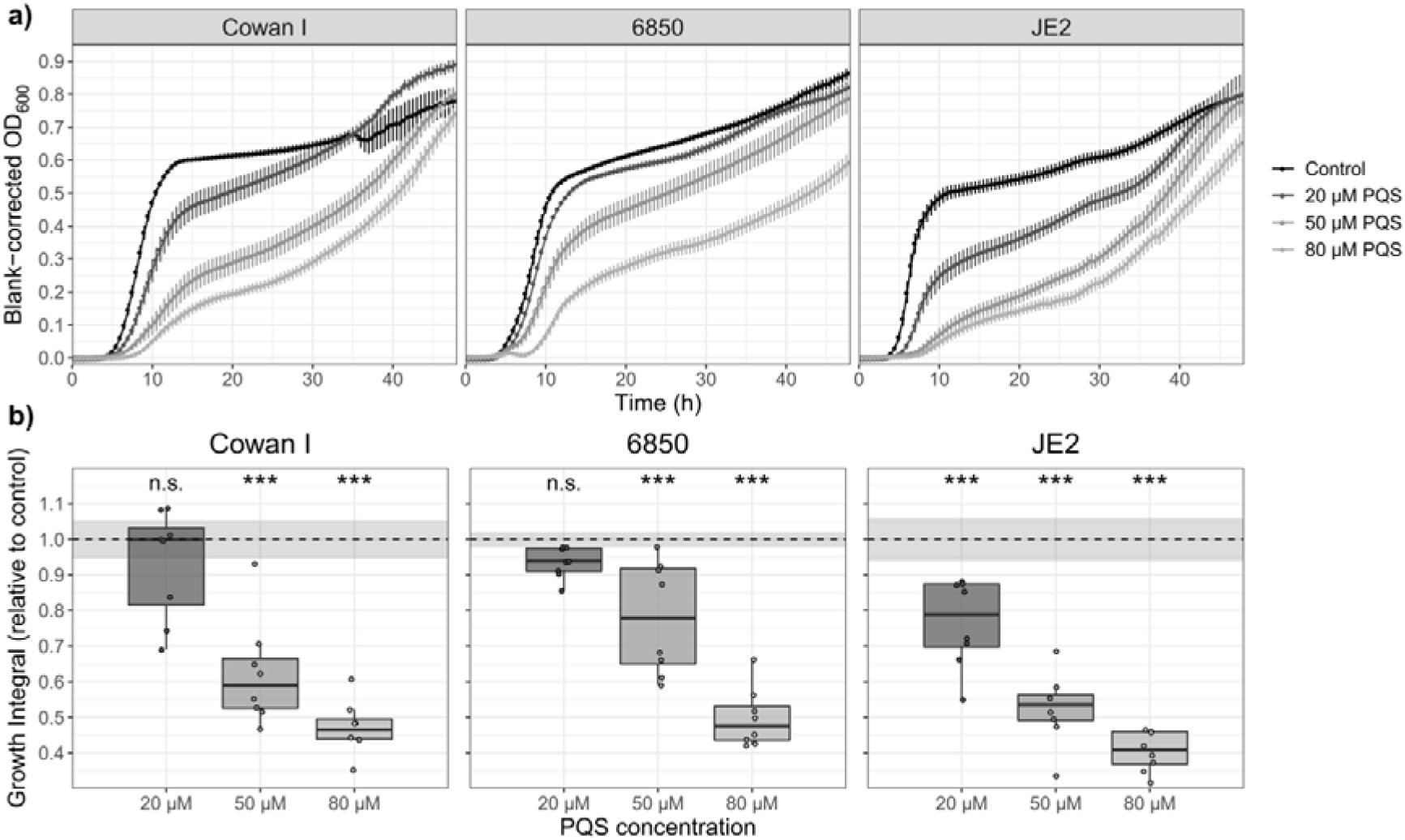
Synthetic PQS suppresses the growth of the three SA strains Cowan I, 6850 and JE2. (a) Growth trajectories of the three SA strains in the presence of increasing concentrations of PQS (20 μM, 50 μM, 80 μM) and TSB containing the same volume of DMSO as the 80 μM PQS condition as control. Shown are the means ± standard errors. (b) Integrals over the growth trajectories of the three SA strains, relative to the control (dotted line at 1.0 ± 95% confidence interval depicted as shaded area). Box plots show the median (bold line), the first and third quartiles, and the 1.5* inter-quartile range (IQR, whiskers) or the range from the lowest to highest value if all values fall within the 1.5* IQR. Asterisks denote significant differences from the control as determined by Tukey’s HSD. *** p < 0.001; n.s. not significant. Data are from two experiments with four independent replicates each per SA strain and condition.

### All SA strains evolve resistance to PA supernatant inhibition

We then asked whether SA adapts to the PA-induced growth inhibition. This is conceivable because the PA supernatant shows anti-bacterial activity, and one could expect SA to evolve resistance against this activity, in similar ways as pathogens evolve resistance against clinically administered antibiotics. To address this question, we performed an experimental evolution experiment, during which we exposed the three SA strains to either 30% PA supernatant in 70% TSB or to 100% TSB over 30 days, in seven-fold replication (Figure S4). Every 48 hours, we transferred evolving cultures to fresh medium using a 1:10^4^ dilution.

Following experimental evolution, we found that populations that had evolved in the presence of PA supernatant showed significantly improved growth performance compared to their respective ancestors when exposed to the PA supernatant, indicating the evolution of resistance (Figure 3a, Cowan I: t_12_ = 11.41, p < 0.0001; 6850: t_12_ = 6.31, p = 0.0002; JE2: t_12_ = 3.61, p = 0.0072). Conversely, all populations that evolved in 100% TSB were still fully or even more susceptible to the PA supernatant compared to the ancestor (Figure 3a, Cowan I: t_12_ = −3.98, p = 0.0055; 6850: t_12_ = 1.00, p = 0.4039; JE2: t_12_ = −0.87, p = 0.4368). Next, we tested whether evolved populations showed TSB-specific adaptations. In support of this view, we found that populations of Cowan I and JE2 that evolved in 100% TSB significantly increased their growth in this medium, but not 6850 (Figure 3b, Cowan I: t_12_ = 11.09, p < 0.0001; 6850: t_12_ = −0.77, p = 0.4590; JE2: t_12_ = 3.70, p = 0.0072). In contrast, populations of all three SA strains that evolved in the presence of PA supernatant did not improve growth in TSB (Figure 3b, Cowan I: t_12_ = 2.32, p = 0.0663; 6850: t_12_ = 1.41, p = 0.2467; JE2: t_12_ = 1.94, p = 0.1137). These results show that all three SA strains have specifically adapted to the presence of PA supernatant and not merely to TSB, and that these adaptations cancel the growth inhibition originally imposed by the PA supernatant.

**Figure 3.**
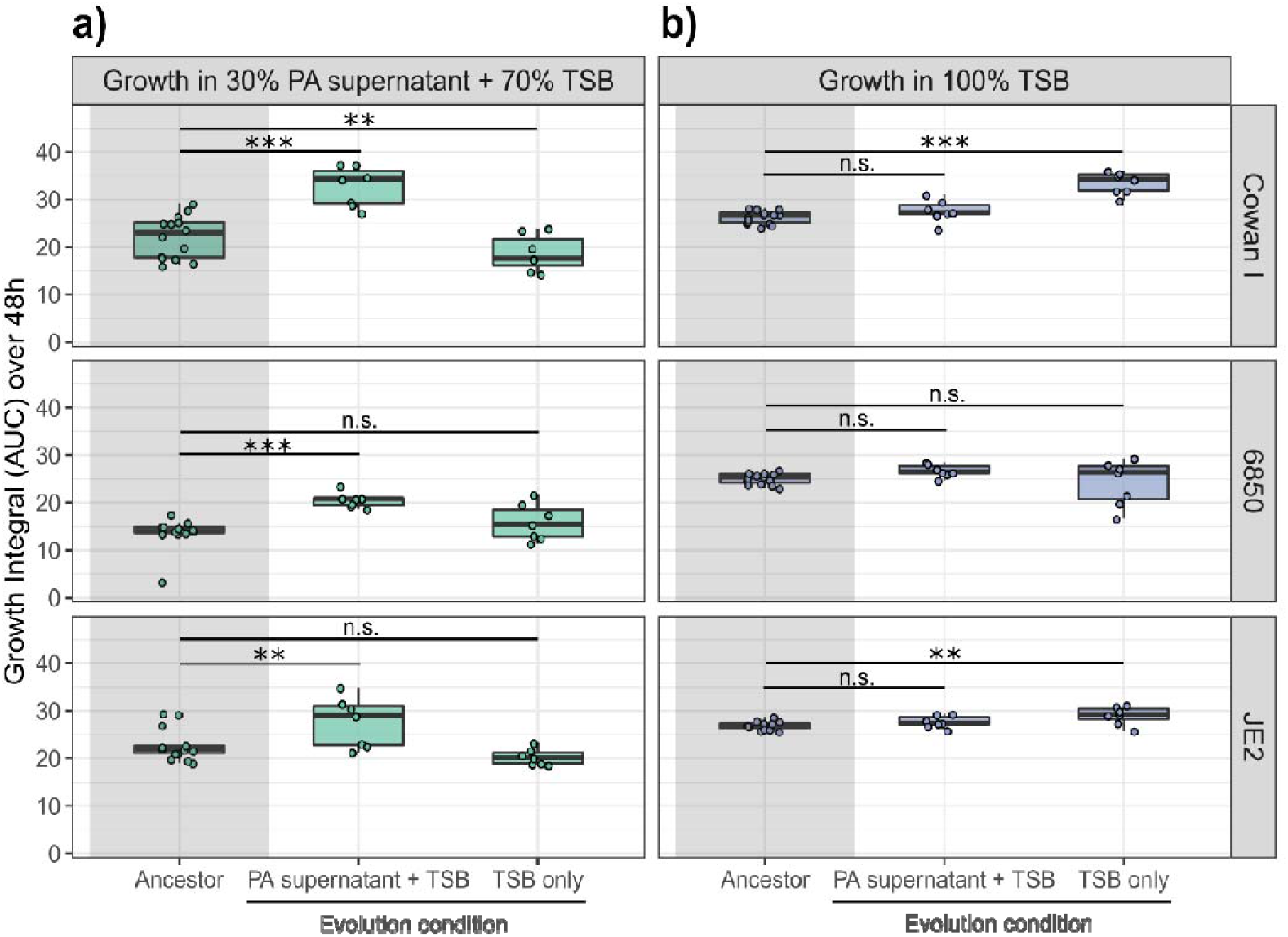
The three SA strains (Cowan I, 6850 and JE2) evolved resistance to inhibitory molecules produced by PA when exposed to its supernatant during experimental evolution. Populations of SA strains (n = 7 per strain) either evolved in 30% PA supernatant + 70% TSB or 100% TSB over 30 days with transfer to fresh medium every 48 hours. (a) Growth of ancestral (grey-shaded area) and evolved populations in medium containing 30% PA supernatant + 70% TSB. SA populations evolved in the presence of PA supernatant significantly improved growth, while populations evolved in TSB alone are still inhibited by the supernatant. (b) Growth of ancestral (grey-shaded area) and evolved populations in 100% TSB medium. SA populations evolved in the presence of PA supernatant did not improve growth in TSB alone, while populations evolved in TSB did so for two of the three SA strains. Box plots show the median (bold line), the first and third quartiles, and the 1.5* inter-quartile range (IQR, whiskers) or the range from the lowest to highest value if all values fall within the 1.5* IQR. Dots represent the individual populations (mean across four technical replicates). Asterisks denote significant differences between the ancestral and evolved populations. *** p < 0.001; ** p < 0.01; n.s. not significant. P-values are adjusted by the false discovery rate method.

We then isolated five random clones per independently evolved lineage (210 clones in total) and repeated the above growth assays with the evolved clones. The results confirmed our population-level analysis of resistance evolution (Figure S5). However, we also observed considerable variation among clones in their growth phenotype, suggesting that populations are heterogenous and consist of multiple different genotypes.

### Clones evolved in PA supernatant show strain-specific resistance to PQS

We then asked whether the increased growth performance of SA clones evolved in PA supernatant is based on specific resistance to the initially inhibitory PQS. To address this question, we exposed evolved clones to 50 μM PQS, representing the intermediate inhibitory concentration in Figure 2. Note that we restricted the above and all subsequent phenotypic screens to 150 clones from 30 out of the 42 populations, as this sample size matched our genome sequencing contingent. We found that the relative growth of evolved SA clones under PQS exposure depended on a significant interaction between strain background and the medium the strains evolved in (ANOVA: F_2,144_ = 4.86, p = 0.0091, Figure 4a). The interaction is driven by 6850 for which clones evolved in PA supernatant grew significantly better under PQS exposure than clones evolved in TSB alone (Tukey’s HSD, p_adj_ < 0.0001). 6850 clones evolved in PA supernatant showed full growth recovery and were on average no longer inhibited by PQS (no significant difference to the growth in TSB alone, t-test: t_49_ = −1.74, p = 0.0883). No such media specific effects were observed for Cowan I (p_adj_ = 0.2519) and JE2 clones (p_adj_ = 0.5475). Although for Cowan I, there were several clones evolved in PA supernatant that showed full growth recovery under PQS exposure. These analyses reveal strain-specific adaptation with 6850 being the only strain where clones showed consistent resistance against PQS.

**Figure 4.**
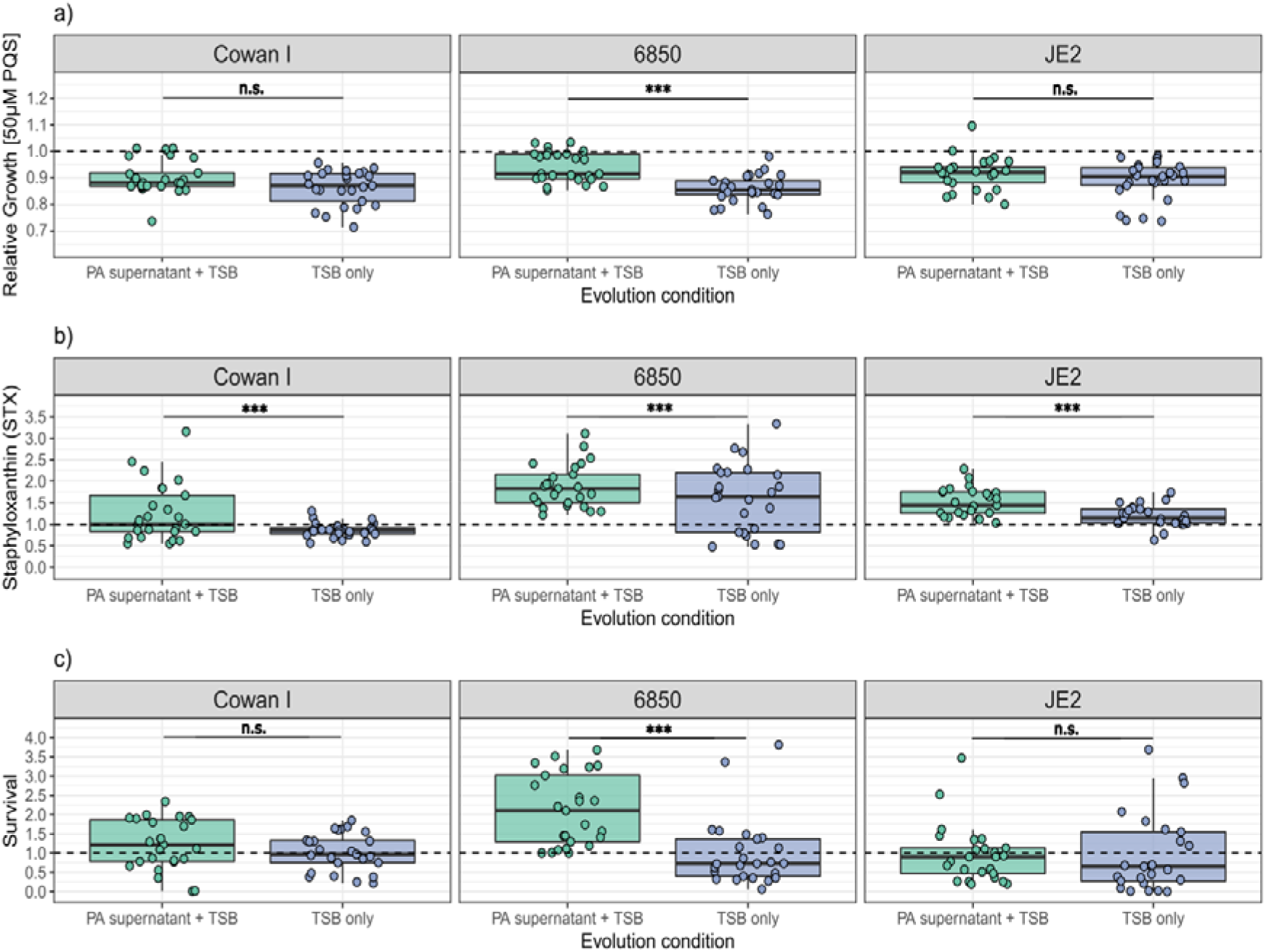
Change in growth under PQS exposure, staphyloxanthin (STX) production and survival in the presence of hydrogen peroxide (H_2_O_2_) after experimental evolution varies across SA strains and media conditions. (a) Growth of evolved clones of Cowan I, 6850 and JE2 when exposed to 50 μM PQS, expressed relative to the untreated respective ancestor (dashed line). (b) Production of STX for evolved clones of Cowan I, 6850 and JE2, expressed relative to the respective ancestor (dashed line). (c) Survival of evolved clones of Cowan I, 6850 and JE2 after exposure for one hour to 1.5% H_2_O_2_, with the survival data being log(x+1)-transformed and expressed relative to the respective ancestor (dashed line). Every dot represents an evolved clone (n=25 for each combination of SA strain and evolution condition) and shows the average value from two independent experiments. Box plots show the median (bold line), the first and third quartiles, and the 1.5* inter-quartile range (IQR, whiskers) or the range from the lowest to highest value if all values fall within the 1.5* IQR. Asterisks denote significant differences between the two evolution conditions (Tukey’s HSD with adjusted p-values). *** p < 0.001; n.s. not significant.

### Clones evolved in the presence of PA supernatant overproduce staphyloxanthin (STX)

PQS and compounds regulatorily controlled by PQS (HHQ and HQNO) promote the formation of reactive oxygen species and therefore cause oxidative stress in target cells (35–38). Since staphyloxanthin (STX), the golden carotenoid pigment, acts as an antioxidant in SA (39), we tested whether the production of this pigment has changed over evolutionary time in the presence of PA supernatant (40). Indeed, we found that STX production was significantly higher in clones evolved in PA supernatant than in clones evolved in TSB alone (ANOVA: F_1,146_ = 16.37, p < 0.0001). Moreover, there were significant differences between SA strains (F_2,146_ = 22.93, p < 0.0001), with the strongest average increase in STX production occurring among the clones of 6850. By contrast, the average increase in STX production was more moderate among Cowan I and JE2 clones, and driven by a few clones in the case of Cowan I. In TSB alone, several evolved 6850 clones also showed increased STX production relative to the ancestor. These analyses suggest that STX production is generally under selection in TSB but reaches particularly high levels in the presence of PA supernatant.

### Survival under oxidative stress correlate with PQS resistance and STX production

The above data supports the scenario that SA can become resistant to oxidative stress induced by PQS and other compounds secreted by PA, and that upregulation of the antioxidant STX might be involved in this process. To test this hypothesis more directly, we exposed all evolved clones for one hour to a 1.5% hydrogen peroxide (H_2_O_2_) solution and explored whether clones evolved in PA supernatant show increased survival under oxidative stress. We observed that the survival of evolved SA clones under H_2_O_2_ exposure depended on a significant interaction between strain background and the medium the strains evolved in (ANOVA: F_2,144_ = 6.81, p = 0.0015, Figure 4b). The interaction is once more driven by strain 6850, for which we found significantly higher survival for clones evolved in PA supernatant than for clones evolved in TSB alone (Tukey’s HSD, p_adj_ < 0.0001). No such media specific effects were observed for Cowan I (p_adj_ = 0.949) and JE2 clones (p_adj_ = 1.0).

Next, we tested for associations between survival in H_2_O_2_ solution and growth under PQS exposure and STX production. For clones evolved in PA supernatant, we indeed found that survival in the presence of H_2_O_2_ correlated positively with growth under PQS exposure (Pearson’s product-moment correlation, r_73_ = 0.30, p = 0.0092, Figure S6a) and STX production (r_73_ = 0.27, p = 0.0207, Figure S6b). In contrast, such correlations were either negative (survival vs. growth under PQS: r_73_ = −0.37, p = 0.0010, Figure S6a) or absent (r_73_ = −0.02, p = 0.8920, Figure S6b) for clones evolved in TSB alone. In sum, our analyses reveal that coping with oxidative stress is a major component of SA adaptation to inhibitory PA supernatant and that the evolutionary response is strongest among clones of 6850.

### SA strains follow divergent and medium-specific evolutionary trajectories

To corroborate our notion that the observed phenotypic changes during experimental evolution are strain and media specific, we performed a principal component analysis (PCA) with the following five phenotypes of the 150 evolved clones: (i) growth in 100% TSB; (ii) growth in 30% PA supernatant + 70% TSB; (iii) relative growth under PQS exposure; (iv) STX production; and (v) survival in the presence of H_2_O_2_. Supporting our notion, we found that evolved clones significantly clustered based on the evolution condition (growth in PA supernatant vs. TSB alone, PERMANOVA; F_1,144_ = 33.24, p = 0.0010, Figure 5), strain background (F_2,144_ = 18.17, p = 0.0010), and their interaction (F_2,144_ = 9.37, p = 0.0010).

**Figure 5.**
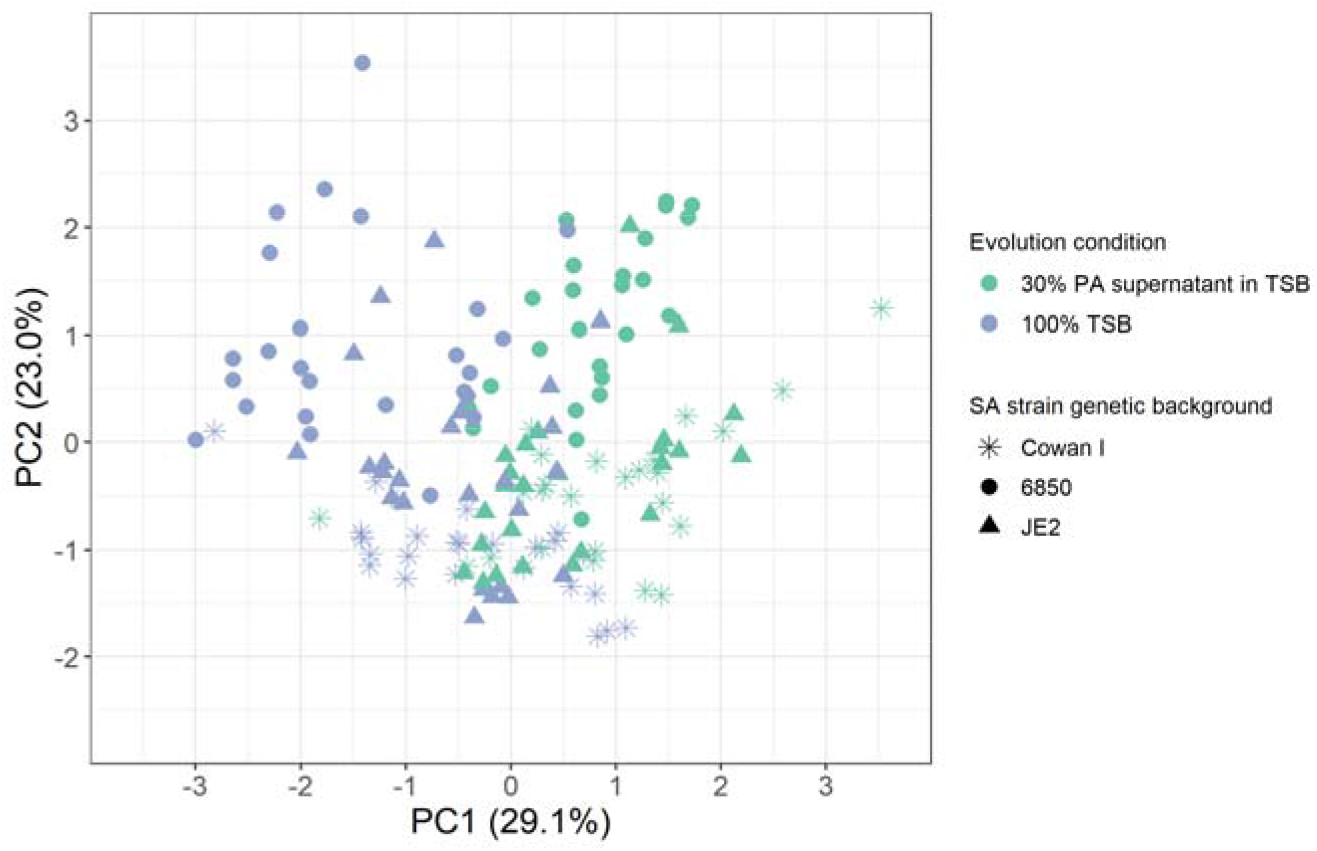
Principal component analysis (PCA) on five phenotypes of 150 evolved clones reveals strain and media specific evolutionary divergence. The following five phenotypes were integrated into the PCA: growth in 30% PA supernatant + 70% TSB, growth in 100% TSB, growth under PQS exposure, staphyloxanthin production and survival in the presence of oxidative stress. The first two principal components (PC1 and PC2) explain 52.1% of the total variance in the data set. Statistical analyses reveal significant divergences in the evolved phenotypes between the three SA strains (Cowan, 6850, JE2), and the condition they evolved in (30% PA supernatant + 70% TSB, 100% TSB).

### Formation of small colony variants in the presence of PA supernatant

Apart from STX, small colony variant (SCV) phenotypes can also offer protection from stressful conditions, including reactive oxygen species (24, 41–43). Therefore, we screened evolved populations and clones for SCV phenotypes. Our first screen involved the plating of evolved populations on agar plates to detect the presence of SCVs. We found SCVs in five out of seven Cowan I populations that had evolved in the presence of PA supernatant (Figure S7). SCVs also surfaced in populations of other strains and media, but at a reduced rate (Cowan I in TSB [2 out of 7 populations]; 6850 in TSB [1/7]; JE2 in PA supernatant + TSB [1/7]). Important to note is that screening for SCVs is challenging, as colony size is a variable trait and phenotypic unstable SCVs can quickly revert back to regular colony sizes (Leimer et al. 2016). Consequently, SCV frequency is typically underestimated.

Nevertheless, the above screen suggests that SCVs arose most commonly in Cowan I populations evolved in PA supernatant. We thus screened all evolved clones from this evolution condition and found that five out of the 25 Cowan I clones expressed a SCV phenotype. We classified three and two of these clones as dynamic/unstable and stable SCVs, respectively. Dynamic/unstable SCVs consistently formed two colony types (small and normal-sized colonies) upon repeated re-streaking. When comparing their growth trajectories in PA supernatant + TSB to the Cowan I ancestor, they had a shorter lag phase and a higher yield (Figure 6a). The two stable SCVs had a longer lag phase and a lower yield than the normal-sized clones isolated from the same population. However, the death phase observed in the Cowan I ancestor was abrogated, suggesting that stable SCV formation increases survival under stressful conditions (Figure 6b). From these analyses, we conclude that both dynamic/unstable and stable SCV formation can be adaptive strategies. While the former increases overall growth performance, the latter prevents cell death in the stationary phase.

**Figure 6.**
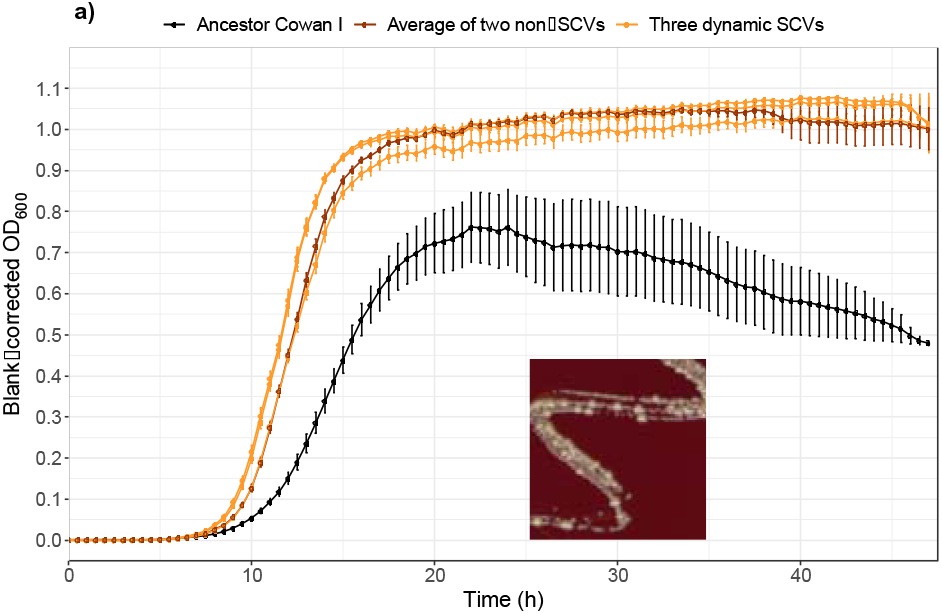
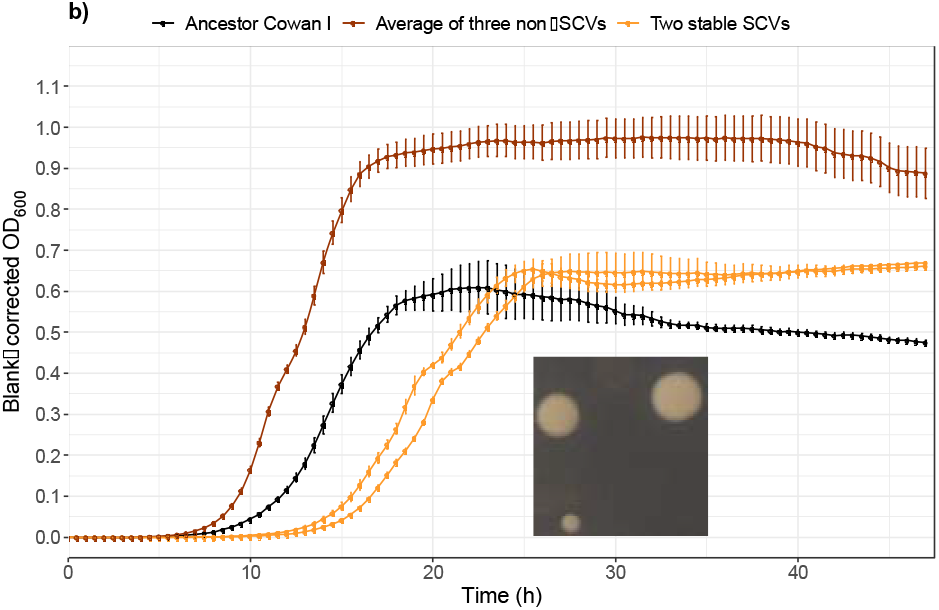
Characteristics of the five small colony variants (SCVs) evolved in the Cowan I background. (a) Growth trajectories of the three dynamic/unstable SCVs in 30% PA supernatant + 70% TSB. Compared to the Cowan I ancestor, dynamic/unstable SCVs have a shorter lag phase and a higher yield, which is similar to the normal-sized clones isolated from the same population. The inset shows one of the dynamic/unstable SCVs with the two colony types on a sheep blood agar plate. (b) Growth trajectories of stable SCVs in 30% PA supernatant + 70% TSB. Compared to normal-sized clones from the same population, the two stable SCVs have a longer lag phase and a lower growth yield, but they no longer show the decrease in OD_600_ that is characteristic for the Cowan I ancestor (see Fig. 1). The inset shows a stable SCV (bottom-left) compared to two normal-sized colonies (upper left and right) from the population from which the two stable SCVs were isolated. Curves show the average growth (± standard error) across four independent replicates.

### The presence of PA supernatant maintains hemolysis

Hemolysis (lysis of red blood cells) is an important virulence-related phenotype in SA, and its expression is regulated in complex ways, including the accessory gene regulator (agr) system and other regulatory genes (44). The loss of hemolysis has repeatedly been observed in SA isolates from chronic infections (45, 46).

Here, we asked whether the presence of PA supernatant affects the evolution of this phenotype in 6850 and JE2 (Cowan I is a non-hemolytic SA strain, due to partial agr dysfunctionality). Our population (Figure S8) and clonal (Figure 7) level analyses show that the majority of 6850 and JE2 clones and populations remained hemolytic after evolution in PA supernatant + TSB. In contrast, after evolution in 100% TSB, hemolytic activity only remained high in clones from 6850 (72% still showed ancestral activity, Figure 7a), while 84% of the JE2 clones lost their hemolytic activity (Figure 7b), a pattern that was also reflected at the population level (Figure S8). These findings show that the presence of PA supernatant fosters the maintenance of hemolysis, and that the speed by which hemolysis can be lost depends on the SA strain genetic background.

**Figure 7.**
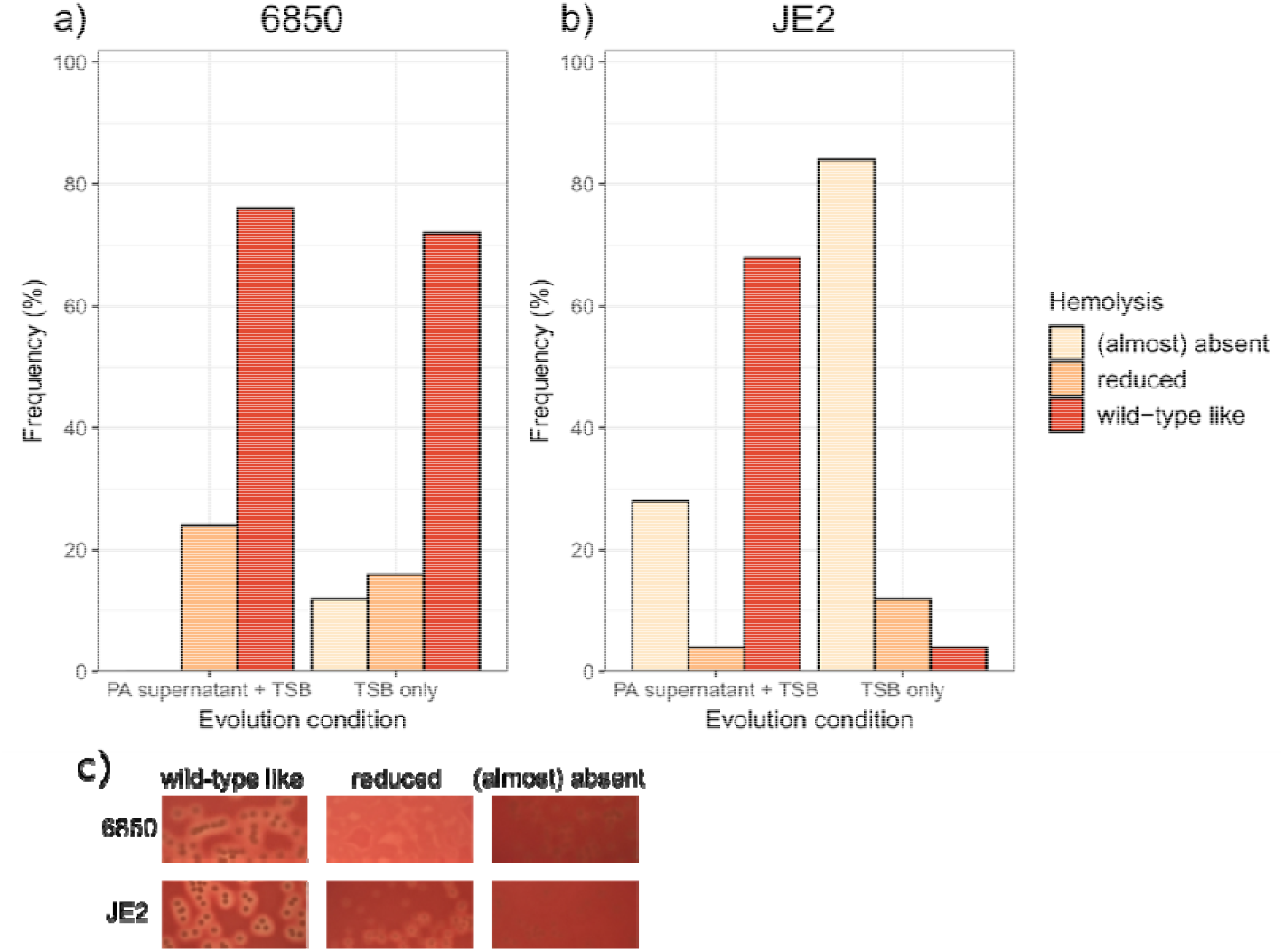
The PA supernatant selects for the maintenance of hemolysis in JE2. 100 evolved clones from the SA strains 6850 and JE2 were plated on sheep blood agar plates to score their hemolysis levels according to three categories: wild-type like (ancestral), reduced, (almost) absent. Note that the Cowan I ancestor is non-hemolytic and therefore not part of this assay. (a) Frequency of the three hemolysis categories for the evolved 6850 clones. There is no significant difference in the hemolysis profiles between clones evolved in PA supernatant + TSB versus in TSB only (Fisher’s exact test: p = 0.2266) and most clones remained hemolytic. (b) Frequency of the three hemolysis categories for the evolved JE2 clones. There is a significant difference in the hemolysis profiles between clones evolved in PA supernatant + TSB versus in TSB only (Fisher’s exact test: p < 0.0001). The large majority of JE2 clones evolved in TSB only are no longer hemolytic. c) Representative pictures of the three hemolysis categories for the SA strains 6850 and JE2.

### Mutational patterns of evolved clones are media-specific

To identify the genetic basis of the observed phenotypic changes, we sequenced the genome of all the characterized 150 evolved clones, and the three SA ancestors Cowan I, 6850 and JE2. Compared to the ancestors, we identified nonsynonymous mutations in 119 different genes and mutations in 51 different intergenic regions across all evolved clones. Among those, 68 and 109 genes or intergenic regions were mutated in clones that had evolved in PA supernatant + TSB and in 100% TSB, respectively. The large majority (96.6%) of the 119 genes with nonsynonymous mutations occurred uniquely in one of the two media (either in PA supernatant + TSB or 100% TSB), and there was little overlap in mutational targets between strains (Figure 8a). These results are in line with our phenotypic assays, highlighting that each SA strain followed a divergent evolutionary trajectory in each of the two media.

**Figure 8.**
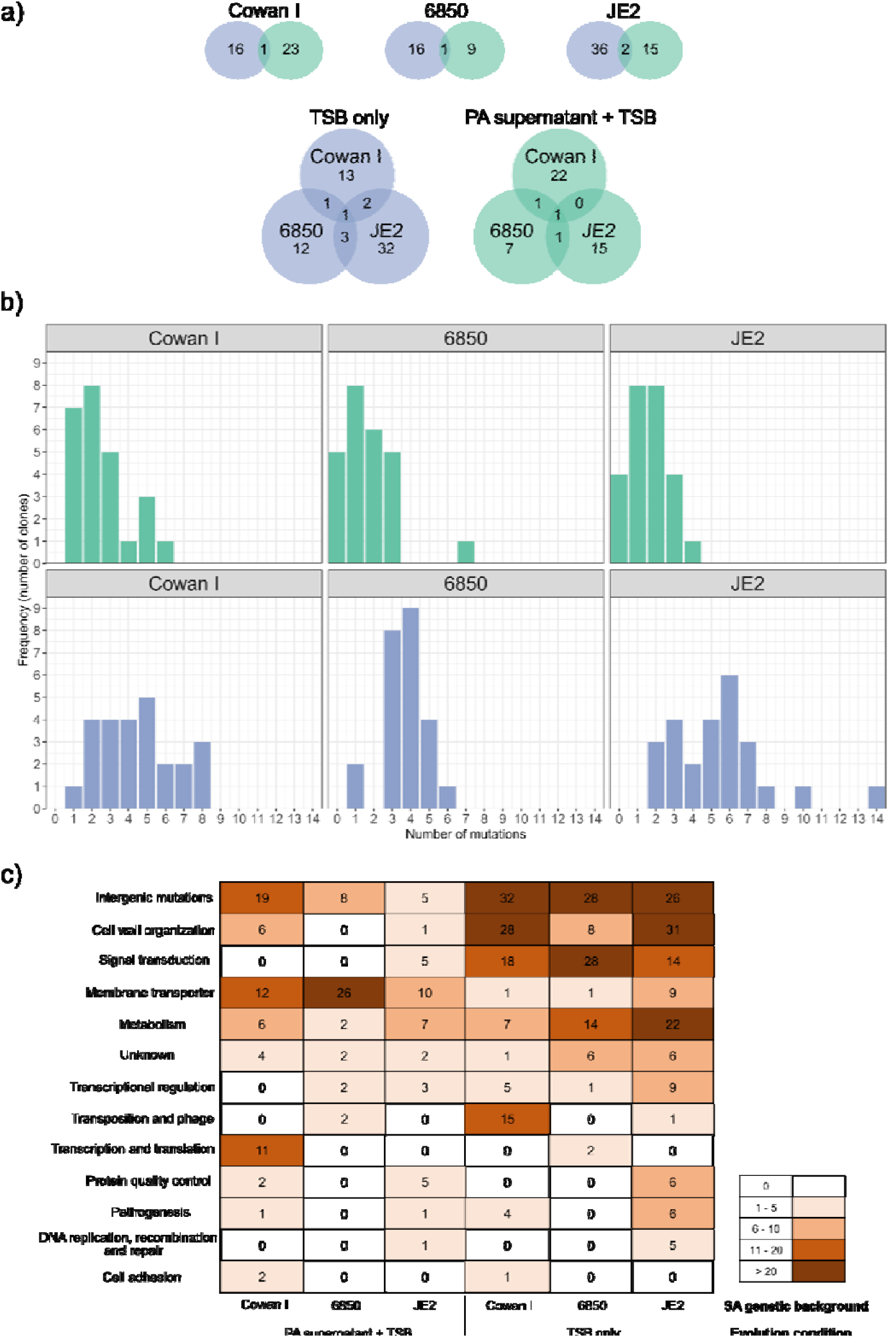
Genomic analysis of 150 evolved clones reveals strain and media specific mutational patterns. a) Venn diagrams showing how many genes with nonsynonymous mutations are unique or shared between evolution conditions (green: 30% PA supernatant + 70% TSB; blue: 100% TSB) and the three SA ancestors (Cowan I, 6850, JE2). b) Number of mutations per clone (nonsynonymous and intergenic mutations) split according to SA strain background and evolution condition. Clones evolved in the presence of PA supernatant had significantly fewer mutations than clones evolved in TSB alone. c) Heatmap showing the total abundance of mutations in functional gene categories and intergenic regions (top row) across all sequenced clones, split by SA strain background and evolution condition. Note that we counted every mutation in every clone for this latter analysis.

For all three SA strains, the median number of mutations was lower for clones evolved in PA supernatant than in TSB alone (Cowan I: 2.0 vs. 4.0; 6850: 1.0 vs. 4.0; JE2: 2.0 vs. 5.0, Figure 8b), indicating that fewer mutations were beneficial in the presence of PA supernatant than in TSB alone. The ratio of nonsynonymous to synonymous SNPs (single nucleotide polymorphism; dN/dS) was higher than one for all strains and in both media (dN/dS for Cowan I: PA supernatant + TSB / 100% TSB = 3.1 / 2.5; for 6850: 13.0 / 12.0; for JE2 3.6 / 4.0). These results show that positive selection and adaptive evolution occurred in both conditions.

To determine whether the mutational patterns differed between the two media and the three SA strains in terms of the functional classes affected, we sorted the mutated genes using aureowiki (Fuchs et al. 2018) and the primary literature. In clones evolved in PA supernatant, we found an enrichment of mutated genes belonging to the category ‘membrane transporter’, while for clones evolved in TSB alone, most mutated genes belonged to the categories ‘metabolism’, ‘cell wall organization’ and ‘signal transduction’ (Figure 8c).

### Parallel evolution in TSB, diverse non-parallel evolution in PA supernatant

Next, we explored whether there is evidence for parallel evolution, which should manifest in similar mutational patterns surfacing across independently evolved populations and strains. For this purpose, we identified genes that mutated more than once in at least two populations/strains. Across both evolution conditions, we found 11 genes that fulfilled these criteria (Table S2). When focusing on clones evolved in TSB alone, there were three genes (*fmtA*, *gdpP*, *walK*) that stood out in terms of both their frequency of mutations and distribution across SA strains. The teichoic acid D-Ala esterase *fmtA* mutated in five Cowan I populations, one 6850 population, and five JE2 populations [5/1/5]. Mutations in the cyclic-di-AMP phosphodiesterase *gdpP* [0/4/5] and the cell wall histidine kinase *walK* [5/3/0] were similarly frequent. A striking pattern is that mutations in any of these three genes are only common in two out of the three SA strains in changing combinations. These results demonstrate that (i) there is high level of parallel evolution between populations of the same strain; (ii) there is a certain level of parallel evolution across strains but the combination of genes that are under selection varies; and (iii) mutations in *fmtA*, *gdpP* and *walK* are likely involved in general adaptations to the TSB medium, as mutations in these genes are absent in clones evolved in PA supernatant.

We found little evidence for parallel evolution among populations evolved in PA supernatant. Only the gene *alsT* mutated in all SA strain backgrounds (Cowan I [2 populations], 6850 [3], JE2 [1]), with *alsT* mutations occurring in a total of 22 clones. This suggests that mutations in *alsT* encoding a glutamine transporter (47), are generally advantageous in the presence of PA supernatant. The lack of any other evidence for parallel evolution indicates that SA populations of all strains adapted to PA supernatant in a diverse manner.

### SA takes diverse mutational trajectories to evolve resistance to PA inhibitory molecules

We used the uncovered mutational patters to construct evolutionary cladograms for each population independently evolved in PA supernatant. Additionally, we mapped three phenotypes (growth in PA supernatant + TSB, staphyloxanthin [STX] production, hydrogen peroxide [H_2_O_2_] survival, the latter correlating with PQS resistance) onto the cladograms to identify links between genotypic and phenotypic changes (Figure 9).

**Figure 9.**
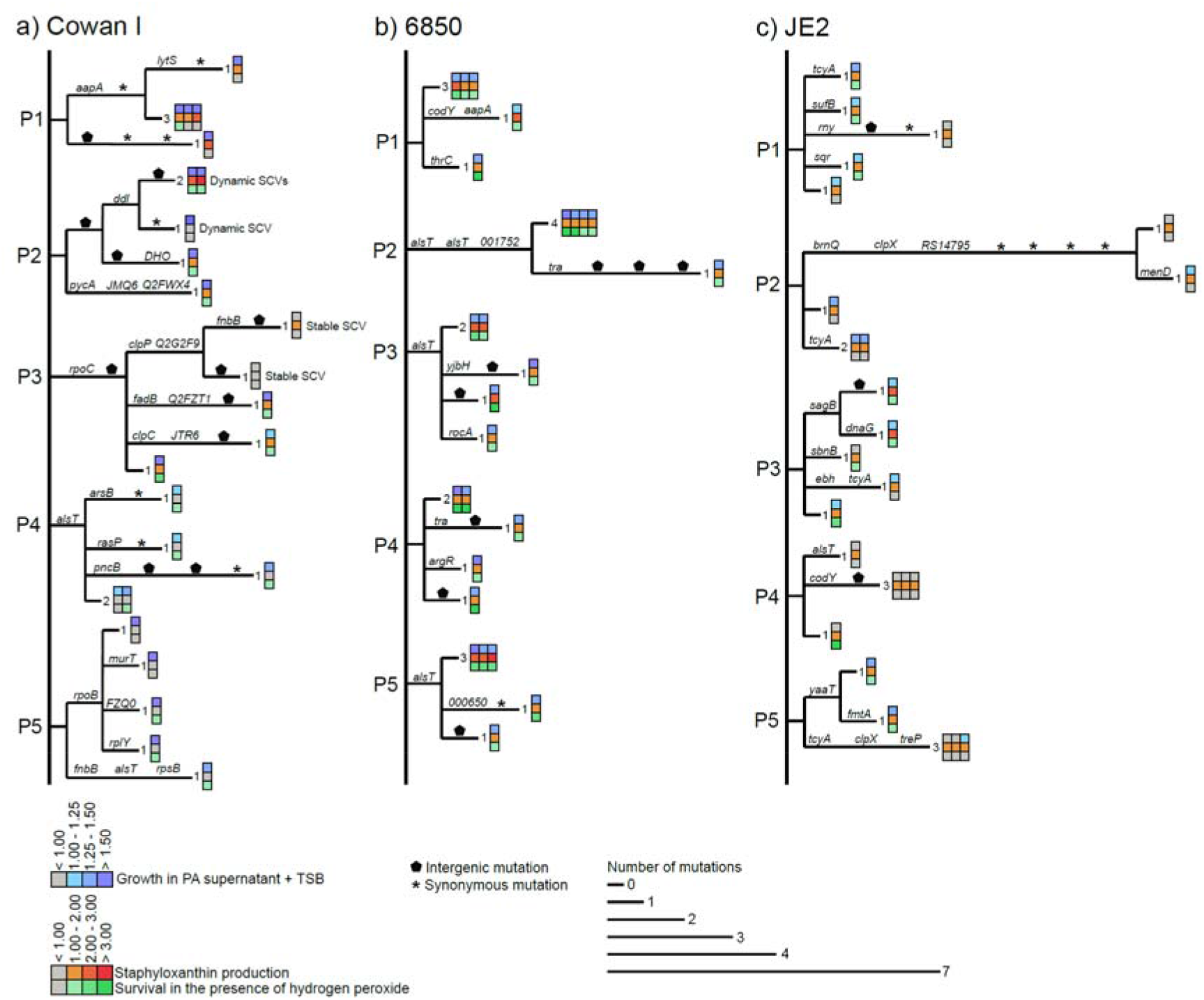
Cladograms showing within-population mutational patterns and their match to phenotypes of SA clones evolved in the presence of PA supernatant. Separate cladograms are shown for each of the five populations (P1 – P5) featuring 25 clones of the three SA strains (a) Cowan I, (b) 6850, and (c) JE2. Branch length corresponds to the number of mutations per clone. Labels on branches indicate either nonsynonymous mutations (gene name or abbreviated locus tag), intergenic mutations (pentagon), or synonymous mutations (asterisk). Synonymous mutations were used to resolve branching patterns among clones. Heatmaps on the tip of the branches show growth in the presence of PA supernatant (top row), staphyloxanthin production (middle row), and survival in the presence of hydrogen peroxide (bottom row) for the respective evolved clones, all expressed relative to the ancestor.

For Cowan I (Figure 9a), mutational patterns were highly diverse within and across populations and there was no apparent link to any of the phenotypes. For example, while 23 out of 25 clones showed resistance to PA supernatant in terms of growth, the underlying mutational patterns were extremely diverse.

For 6850 (Figure 9b), there is some evidence for parallel evolution via mutations in *alsT*, which became fixed in three different populations. All clones with *alsT* mutations showed higher growth, increased STX production and H_2_O_2_ survival. However, the same phenotypic changes were also observed in other 6850 clones without *alsT* mutations. This shows that mutations in different genes can lead to the same phenotypic changes.

For JE2 (Figure 9c), the genotype-phenotype matching was diverse, too. Overall, JE2 showed the least pronounced changes in phenotypes, most likely because the ancestor was least affected by the PA supernatant (Figure 1). But when focusing on the clones in populations P1, P3 and P5, which showed the most pronounced increase in growth, STX production and H_2_O_2_, we found completely different underlying mutational patterns in each population. For example, mutations in *tcyA* (population P1), *sagB* (P3), and *yaaT* (P5) all seemed to be associated with the altered phenotypes. The *tcyA* gene, encoding a membrane transporter, stood out from the other genes as it was repeatedly mutated in seven clones from four different JE2 populations, suggesting that it could play a more general role in SA resistance evolution.

## Discussion

*Pseudomonas aeruginosa* (PA) and *Staphylococcus aureus* (SA) are among the most troublesome pathogens in polymicrobial infections, such as wound and cystic fibrosis (CF) lung infections. That is the reason why there is high interest in understanding how species interactions shape disease outcome (3, 9, 48, 49). A large body of literature has examined PA-SA interactions at the molecular level, and mostly described PA as the dominant species suppressing the growth of SA by producing a variety of inhibitory molecules (8). However, polymicrobial infections containing PA and SA are often chronic so that the two species may adapt to one another, and SA may become resistant to PA inhibitory molecules (50, 51). Here, we tested this hypothesis by exposing three different SA strains (Cowan I, 6850 and JE2) to the growth-inhibitory supernatant of the PA reference strain PAO1 during experimental evolution. We found that all SA strains rapidly evolved resistance to the PA induced growth inhibition. Using a combination of phenotypic and genotypic assays, we show that (i) resistance was associated with the upregulation of traits involved in the protection from reactive oxygen, including staphyloxanthin (STX) production and the formation of small colony variants (SCV), (ii) adaptive patterns were SA strain-specific, and (iii) mutations in genes encoding membrane transporters were the most frequent target of evolution. Thus, our results suggest that resistance is achieved by a combination of decreased influx and increased efflux of harmful PA compounds, together with direct protective measures against oxidative stress. Since these evolutionary changes concern virulence traits and membrane transporters, the observed SA adaptations to PA supernatant could spur higher virulence and increased antibiotic resistance in infections.

All three SA strains were initially compromised in their growth by the PA supernatant, and we confirmed that the *Pseudomonas* quinolone signal (PQS) molecule was involved in growth inhibition. PQS does not only serve as a quorum-sensing signaling molecule but also functions as an iron trap (52). The iron-chelating activity of PQS reduces the bioavailability of iron in the environment, and this was shown to have an inhibitory effect on a variety of gram-negative and gram-positive bacterial species (32, 53). While the induction of iron limitation could have contributed to the observed growth inhibition in our experiments, our data rather suggest that inhibition was mainly due to the ability of PQS to trigger the formation of reactive oxygen species (35–38). Furthermore, HQNO is another molecule produced through the PQS synthesis cascade that also has anti-SA properties via inhibition of the respiratory chain (24). It is thus likely that the inhibitory effects on SA growth observed in our supernatant assays is a combination of the effects of PQS and HQNO.

We found that the extent to which SA strains were inhibited and the strength of resistance evolution towards the PA supernatant varied across SA strains (Figure 3). While the Cowan I ancestor was substantially inhibited, it showed the largest improvement in growth after evolution in the PA supernatant, and even outperformed the ancestral wildtype growing in TSB alone. Similarly, 6850 was initially greatly inhibited by the PA supernatant, and all replicate populations showed resistance evolution, reflected by a significantly improved growth. JE2 was initially the least inhibited strain, and while overall, we found significantly improved growth after evolution, not all replicate populations followed this trend. This differential response matches evolutionary predictions that the most severely affected SA strains face the strongest selection pressure and consequently should show the strongest response in terms of resistance evolution. Regarding JE2, it is interesting to note that this strain is the most virulent and most competitive SA strain in our panel (19, 54). It could well be that it has already interacted with PA in its past and might therefore already be equipped to deal with PA inhibitory molecules, leaving less room for further fitness improvements compared to the less virulent and less competitive Cowan I or 6850 strains.

We observed that SA virulence traits were under selection and often overproduced, which could have consequences in infections (55, 56). Importantly, the virulence traits under selection varied across SA strains, reinforcing our notion that evolutionary responses to PA inhibitory molecules are SA strain specific. Increased formation of SCVs evolved almost exclusively in Cowan I populations. However, we assume that the true SCV frequency might be higher than the one reported, because the frequency of SCVs is typically underestimated. The formation of SA SCVs is a well-known adaptive response towards stressful conditions (43), and we propose that SCVs evolved in our experiment as a response to PA inhibitory molecules such as PQS and HQNO (24). Due to their slow metabolism, SCVs frequently exhibit reduced antibiotic susceptibility, which makes them particularly problematic in a clinical context (57). For 6850 clones, the strongest phenotypic changes involved increased STX production and improved survival under oxidative stress (PQS and H_2_O_2_ exposure). In the context of infections, such adaptations would be problematic, because innate host immune responses against pathogens typically involve the production of reactive oxygen species (58). For JE2, we observed that hemolysis was selectively maintained in the presence of PA supernatant but lost in its absence. While hemolysis is an important virulence trait, it can be lost in chronic infections (45, 46). Our findings now suggest that the presence of PA could maintain this trait and keep JE2 and related strains at a higher state of virulence. At the genetic level, we found positive selection but little evidence for parallel evolution across strains and populations upon exposure to the PA supernatant. This indicates that mutations in many different genes were under selection and possibly contributed to resistance evolution towards PA inhibitory molecules. The fact that mutations in a broad spectrum of genes can promote resistance highlights the enormous evolvability of SA (59). In contrast, we found evidence for parallel evolution among clones that evolved in the TSB medium alone, where we identified several genes (Table S2) that were frequently mutated in independent populations and across strains. These findings show that, whether parallel evolution occurs depends on the environment and the specific selection pressures that prevail therein.

Although there was little evidence for parallel evolution at the gene level among clones evolved in PA supernatant, we found parallelism at the functional level, with many mutations occurring in membrane transport genes (Figure 8c). For example, the gene encoding the glutamine transporter AlsT was mutated in 22 clones from six populations across all three SA strains. While AlsT inactivation reduces the susceptibility of SA to the toxic glutamine analogue γ-L-glutamyl hydrazide (47), our results now suggest that AlsT alterations might similarly reduce the uptake of PA inhibitory molecules. Moreover, it was shown that loss-of-function mutations in *alsT* increase intracellular c-di-AMP levels, which allows bacteria to better resist stressful conditions (60). Our results support this notion and suggest that *alsT* mutations were selected for to withstand the stress imposed by PA inhibitory molecules. Another example involves mutations in the gene *tcyA*, which encodes a part of the cystine transporter TcyABC. While mutations in this gene were unique to JE2, they surfaced in seven clones in four out of the five JE2 populations. Deletion of *tcyA* leads to resistance against the toxic cystine analogue selenocystine in SA (61). While PA PAO1 indeed contains the selenocysteine synthesis cluster, it is currently not known how its expression is controlled and whether selenocystine is deployed against competitors. Our data now suggest that it might indeed play a role in competition, and that mutations in *tcyA* could be an evolutionary response by SA to reduce selenocystine uptake. As membrane transport systems are known to control virulence and antibiotic resistance in SA, they have been proposed as novel targets for therapeutic agents (62). Our finding that the exposure to PA inhibitory molecules can select for SA variants with altered membrane transporters could therefore pose a problem for drug design and therapy, as these mutants could be less susceptible to antibiotics.

In summary, our work shows that SA rapidly and specifically adapts to PA inhibitory molecules. Several virulence traits were upregulated or selectively maintained in response to PA supernatant exposure, and our genomic analysis revealed that mutations in membrane transport genes are associated with resistance evolution in SA. Such adaptations could have severe implications for infections, as SA virulence might be increased, and baseline levels of antibiotic resistance might arise even before a treatment is applied. Thus, a key next step would involve testing whether the observed increases in virulence trait expression and mutations in membrane transporters lead to altered virulence and antibiotic susceptibility profiles *in vivo*. Moreover, while we focused on SA evolution and others on PA evolution (30), we urgently need to study co-evolution to understand the patterns and consequences of reciprocal adaptation in both species, as they might occur in polymicrobial infections (49). In a broader context, the fact that evolutionary trajectories were strongly SA strain-specific suggests that we ultimately need to understand polymicrobial infection dynamics and its consequences at an ‘individualized’ patient-by-patient level in order to improve treatment options.

## Materials and Methods

### Bacterial strains, media, and growth conditions

We used *Pseudomonas aeruginosa* (PA) strain PAO1 (ATCC 15692) and the three genetically different *Staphylococcus aureus* (SA) strains Cowan I, 6850 and JE2 (Table S1, see also (19, 63) for further strain information and species interaction patterns) for all our experiments. All growth experiments and the experimental evolution were performed in tryptic soy broth (TSB, Becton Dickinson, Heidelberg, Germany) medium. Overnight cultures (prior to experimental evolution) were grown in 10 ml TSB in 50 ml falcon tubes at 37 °C and 220 rpm with aeration. The phenotypic assays after experimental evolution involved large sample sizes including many evolved clones, populations, and the respective ancestors. For this reason, we grew overnights in 24-well plates filled with 1.5 ml TSB per well at 37 °C and 170 rpm.

### Supernatant assays

To test whether the supernatant of PA inhibits SA (and vice versa) we performed supernatant growth assays as follows. We first generated cell-free supernatants by growing PA and SA donor strains overnight in TSB. After centrifugation of the cultures to pellet cells, we sterile-filtered the supernatant using 0.2 μm pore size filters (Whatman, Fisher Scientific, Reinach, Switzerland) and stored aliquots of the supernatant at −20 °C. For the supernatant assay, we grew PA and SA receiver strains overnight, washed bacterial cell pellets with 10 ml 0.8% NaCl and adjusted OD_600_ (optical density at 600 nm) to obtain comparable cell numbers per ml for all strains. To achieve this, we adjusted OD_600_ of PA to 1.0, for SA strains JE2 and 6850 to 0.4 and for Cowan I to 0.8. Subsequently, we diluted the PA inoculum 1:10^5^ and the SA inoculums 1:10^4^ to start our experiments. PA was diluted more because it grows faster compared to SA. The diluted inoculums were prepared in the following three media conditions: (1) 100% TSB (as growth control); (2) 30% NaCl solution + 70% TSB (control to mimic reduced nutrient availability); (3) 30% PA or SA supernatant + 70% TSB (condition of interest).

Supernatant assays were performed in 96-well plates with 200 μl per well. We incubated plates statically at 37 °C in a plate reader (Tecan Infinite M nano, Tecan, Männedorf, Switzerland) and measured growth of strains by recording OD_600_ every 30 min for 48 hours, with 30 sec shaking events prior to recordings. Effect of PA supernatant on growth of SA strains was assessed in seven independent experiments and effect of SA supernatants on PA growth was assessed in one experiment per SA supernatant. In every assay, we used four replicates per strain and growth condition.

### SA growth in the presence of PQS

To test whether PQS (heptyl-3-hydroxy-4(1H)-quinolone, Sigma-Aldrich, Buchs, Switzerland) is involved in the growth inhibition of SA, we exposed cultures of all three SA strains to final concentrations of 20 μM, 50 μM or 80 μM PQS (10 mM stock in DMSO), supplemented to the TSB growth medium, and compared their growth relative to the growth in TSB supplemented with DMSO alone (same volume of DMSO as for the 80 μM PQS treatment). SA strain preparation and growth measurements (OD_600_) in 200 μl medium volume in 96-well plates followed the exact same protocol as for the supernatant assays using a plate reader.

Following evolution, we repeated the above experiment for 150 evolved clones (see below for the selection procedure) to test whether SA strains have evolved resistance to the growth inhibition induced by PQS. For this experiment, we exposed each individual clone (grown overnight in TSB) to either 50 μM PQS or DMSO (as a control) in triplicates. We followed the same protocol as above, except that we incubated the plates in a shaken incubator at 170 rpm and measured growth at the final time point after 48 hours.

### Experimental evolution of SA in the presence or absence of PA supernatant

To elucidate whether and how SA strains respond to the presence of PA supernatant over time, we performed an experimental evolution experiment, in which we exposed the three SA strains Cowan I, 6850 and JE2 for 30 days to either 30% PA supernatant + 70% TSB (experimental treatment) or 100% TSB (control treatment). Experimental evolution started from clonal populations. We had seven independent populations per strain and condition, resulting in a total of 42 evolving populations, distributed on 96-well plates (one plate per SA ancestor). All populations were incubated statically at 37 °C. Every 48 hours, we diluted and transferred the evolving cultures 1:10,000 into fresh medium and recorded OD_600_ as a proxy for growth prior to each transfer using a plate reader (Tecan Infinite M nano). To the part of the culture that was not transferred, we added equal volumes of a sterile 85% glycerol solution directly into the wells of the 96-well plate and froze them at −80 °C as a backup. Overall, the experiment ran for 30 days (15 transfers). Based on a pre-experiment, we estimated the mean number of doublings (± standard deviation) for the three SA ancestors in the two growth conditions during a 48-hour growth cycle as follows: for Cowan I: 14.5 ± 0.4 (in TSB) / 9.2 ± 0.7 (in PA supernatant + TSB); for 6850: 14.3 ± 0.8 / 10.6 ± 1.7; for JE2: 14.6 ± 0.5 / 13.5 ± 0.6. Thus, our experimental evolution involved at least 140 - 220 bacterial generations (assuming no evolutionary improvements in growth). Following experimental evolution, we plated all evolved populations on 1.2% TSB agar and randomly picked five clones per population for further characterization. All selected clones were first cultured overnight in TSB and then frozen at −80 °C by mixing 50% of culture with 50% of a sterile 85% glycerol solution.

### Growth of evolved populations and clones

We tested whether the 42 evolved populations and the selected 210 clones showed improved growth performance relative to the ancestors in the two conditions of our experimental evolution (30% PA supernatant + 70% TSB and 100% TSB). We exposed all populations and clones to the condition they had evolved in, but also to the alternate condition they had not evolved in. For this growth screening, we followed the exact same protocol as the one described above for the supernatant assay.

### Staphyloxanthin (STX) quantification

For the subsequent phenotypic and genotypic assays, we used a total of 150 clones from 30 populations (5 clones per population). Sample size was reduced to meet the sequencing contingent of this project. In a first assay, we tested whether evolved clones changed staphyloxanthin (STX) production compared to the respective ancestor and quantified STX based on a previously published protocol (40). Briefly, we first collected 1 ml from each culture of all evolved clones and ancestors (grown overnight in TSB) and resuspended washed pellets in 400 μl methanol. We incubated the samples for 10 min at 55 °C, pelleted cells by centrifugation and collected the supernatant containing the STX. Absorbance of the crude extract was measured in duplicates at λ = 465 nm using a plate reader (Tecan Infinite M200 pro). We blank-corrected STX values and expressed them first, relative to the blank-corrected OD_600_ of the respective clone and second, relative to the respective STX value of the ancestor.

### Hydrogen peroxide (H2O2) survival assay

To determine whether evolved clones are better at surviving under oxidative stress conditions than the respective ancestors, we performed a hydrogen peroxide (H_2_O_2_) survival assay using a previously published protocol (64). In short, we subjected cultures of all evolved clones and ancestors (grown overnight in TSB) to either 1.5% H_2_O_2_ or 0.8% NaCl (as control) for one hour at 37 °C. Subsequently, we plated appropriate dilutions on 1.2% TSB agar, incubated plates overnight at 37 °C and enumerated colony forming units (CFUs) on the next day. We then calculated the percentage survival for each clone as the CFUs from the H_2_O_2_ treatment divided by the CFUs from the control treatment multiplied by 100. Finally, we expressed survival relative to the respective ancestor.

### Small colony variant (SCV) detection and quantification

To determine whether evolved clones express SCV phenotypes, we plated cultures of all evolved clones on 1.2% TSB agar, incubated plates overnight at 37 °C, and then classified the colonies with an area smaller than 20% of the ancestral colony size as SCV. To estimate the prevalence of SCVs at the population level, we also serially diluted and plated aliquots of every evolved population on 1.2% TSB agar and assessed the percentage of SCVs per population after incubation of plates overnight at 37 °C.

### Hemolysis assay

The ancestral strains of 6850 and JE2 are known for their ability to lyse red blood cells (hemolysis), whereas Cowan I is unable to do so. To determine whether evolved clones of 6850 and JE2 show different hemolysis patterns compared to their ancestors, we plated cultures of all evolved clones and ancestors (grown overnight in TSB) on COS plates (columbia agar + 5% sheep blood, Biomérieux, Petit-Lancy, Switzerland). COS plates were incubated overnight at 37 °C. On the next day, we qualitatively assessed the hemolysis pattern of each clone according to three discrete categories: ancestral-like hemolysis (wild-type like), reduced hemolysis, (almost) absent hemolysis (examples depicted in Figure 6c). In addition, we also plated all evolved populations on COS plates to assess the frequency of the three hemolysis phenotypes within populations.

### Genome sequencing

To identify genetic changes in evolved clones relative to their ancestors, we extracted the genomic DNA of all 150 phenotypically characterized clones and the three ancestral SA strains. We used the Maxwell RSC Cultured Cells DNA Kit (Promega, Dübendorf, Switzerland) together with the Maxwell RSC 48 instrument (Promega) following the manufacturer’s protocol. To lyse the gram-positive SA cells, we added lysostaphin (L7386, Sigma-Aldrich) to the samples (final concentration of 25 μg per 400 μl sample). DNA concentrations were quantified using the QuantiFluor dsDNA sample kit (Promega). Library preparation was performed with 100 ng of DNA input using the TruSeq DNA Nano kit (Illumina, San Diego, USA) according to the manufacturers’ instructions. The libraries were quantified using the Tapestation (Agilent Technologies, Santa Clara, USA) and qPCR (Roche, Rotkreuz, Switzerland) and equimolarly pooled. Finally, the libraries were sequenced 150 bp paired-end on the NovaSeq6000 system (Illumina, San Diego, USA). The samples had an estimated coverage of 100x for the evolved clones and 200x for the three SA ancestors.

The quality of the raw sequencing data was assessed using FastQC (version 0.11.9) (https://www.bioinformatics.babraham.ac.uk/projects/fastqc/) and a contamination check was performed using Fastqscreen (version 0.14.1). Subsequently, the raw reads were preprocessed using Trimmomatic (65) with the following settings: trimmString="ILLUMINACLIP: adapters.fa:1:30:10 LEADING:15 TRAILING:15 SLIDINGWINDOW:5:30 AVGQUAL:32 MINLEN:80”. All the reads that passed the trimming and quality filtering steps were analyzed using Snippy (https://github.com/tseemann/snippy) (v4.5.2) with default parameters. For the ancestral strains 6850 and JE2, the reads were aligned to the published reference genomes (GenBank accession Nr. CP006706.1 and NZ_CP020619.1, respectively). Variants that were present in the ancestor compared to the published reference genome were removed and only those variants that occurred during experimental evolution were kept. Finally, we used SnpEff version 4.3t (built 2017-11-24 10:18) (https://pcingola.github.io/SnpEff/) to predict variant effects. For Cowan I, no suitable reference genome was available, and we thus performed a *de novo* assembly with SPAdes (v3.12.0) and annotation with Prodigal (v2.6.3) using the reads from our ancestor, and then directly called the variants as in the other two strains for the evolved clones relative to the *de novo* assembled ancestral genome. Note that one clone had to be discarded after library preparation due to poor DNA quality.

### Statistical analysis

All statistical analyses were performed with R Studio version 3.6.3. We used general linear models wherever possible and consulted Q-Q plots and the Shapiro-Wilk test to examine whether residuals were normally distributed. The basic linear model that we used was an analysis of variance (ANOVA) in which we put the response variable in relation to the manipulated factors, which are (i) the SA strain background (Cowan I, 6850, JE2), (ii) the experimental conditions (presence/absence of PA supernatant or PQS concentration), and (iii) the interaction between the two. We used variants of this model for statistical analysis of all growth data, STX production, and survival in the presence of H_2_O_2_ as response variables. Non-significant interactions were removed from the models. If data was not normally distributed (as for the survival data), we log_10_(x+1)-transformed all values for statistical analysis and scaled them back to the respective ancestor for plotting.

We performed one sample t-tests to test whether growth under PQS exposure, STX production and survival in the presence of H_2_O_2_ in evolved clones was significantly different from the expected ancestral value. The false discovery rate method was used to correct p-values whenever necessary. For frequency comparisons between hemolysis categories for 6850 and JE2 clones evolved in the two media, we performed Fisher’s exact tests. The principle component analysis (PCA) was performed on clonal phenotypes using the *vegan* package in R. We tested for inference using permutational multivariate analysis of variance (PERMANOVA).

## Competing interests

The authors declare that no competing interests exist.

## Author Contributions

SN and RK designed research, SN and LS performed research, LP and JG analyzed the genome sequencing data, SN, LS and RK analyzed all other datasets and wrote the paper with inputs from LP and JG. All authors approved the manuscript prior to submission.

## Funding

This project has received funding from the European Research Council under the European Union’s Horizon 2020 research and innovation program (grant agreement no. 681295) to RK, from the Swiss National Science Foundation (31003A_182499) to RK, and from the University of Zürich Teaching Fund (2019_12 / Fostering OMICS research through research-based teaching and learning) to LP and JG.

## Acknowledgements

We thank Markus Huemer and Annelies Zinkernagel (University Hospital of Zürich) for providing *S. aureus* strains, the Eberl group (University of Zürich) for providing the *P. aeruginosa* Δ*pqsA* strain, Jay Tracy for helping with DNA extraction, Maria Domenica Moccia for performing the whole genome sequencing, Giancarlo Russo for bioinformatic analysis, and Alexandre Figueiredo for comments on the manuscript. Illustration for Supplementary Figure 4 was created using BioRender (www.biorender.com).

## Data availability statement

All source data associated with this study will be deposited in an online repository upon the acceptance of the manuscript.

## Notes

### Competing Interest Statement

The authors have declared no competing interest.

### Summary of Updates

This version of the manuscript has been revised to update Figure 4, now containing new data.

## References

1. Brogden KA, Guthmiller JM, Taylor CE. 2005. Human polymicrobial infections. The Lancet 365:253–255.

2. Peters BM, Jabra-Rizk MA, O’May GA, Costerton JW, Shirtliff ME. 2012. Polymicrobial interactions: Impact on pathogenesis and human disease. Clin Microbiol Rev 25:193–213.

3. Short FL, Murdoch SL, Ryan RP. 2014. Polybacterial human disease: the ills of social networking. Trends Microbiol 22:508–516.

4. Lim NCS, Lim DKA, Ray M. 2013. Polymicrobial versus monomicrobial keratitis: a retrospective comparative study. Eye Contact Lens 39:348–354.

5. Pammi M, Zhong D, Johnson Y, Revell P, Versalovic J. 2014. Polymicrobial bloodstream infections in the neonatal intensive care unit are associated with increased mortality: a case-control study. BMC Infect Dis 14:390.

6. Tan TL, Kheir MM, Tan DD, Parvizi J. 2016. Polymicrobial periprosthetic joint infections: outcome of treatment and identification of risk factors. J Bone Joint Surg Am 98:2082–2088.

7. Jorge LS, Fucuta PS, L MG, Nakazone MA, de JA, Chueire AG, Costa MJ. 2018. Outcomes and risk factors for polymicrobial posttraumatic osteomyelitis. J Bone Jt Infect 3:20–26.

8. Hotterbeekx A, Kumar-Singh S, Goossens H, Malhotra-Kumar S. 2017. In vivo and In vitro Interactions between *Pseudomonas aeruginosa* and *Staphylococcus spp*. Front Cell Infect Microbiol 7:106.

9. Limoli DH, Hoffman LR. 2019. Help, hinder, hide and harm: What can we learn from the interactions between *Pseudomonas aeruginosa* and *Staphylococcus aureus* during respiratory infections. Thorax 74:684–692.

10. Gjødsbøl K, Christensen JJ, Karlsmark T, Jørgensen B, Klein BM, Krogfelt KA. 2006. Multiple bacterial species reside in chronic wounds: a longitudinal study. Int Wound J 3:225–231.

11. Dowd SE, Sun Y, Secor PR, Rhoads DD, Wolcott BM, James GA, Wolcott RD. 2008. Survey of bacterial diversity in chronic wounds using pyrosequencing, DGGE, and full ribosome shotgun sequencing. BMC Microbiol 8:43.

12. Hubert D, Réglier-Poupet H, Sermet-Gaudelus I, Ferroni A, Bourgeois ML, Burgel PR, Serreau R, Dusser D, Poyart C, Coste J. 2013. Association between *Staphylococcus aureus* alone or combined with P*seudomonas aeruginosa* and the clinical condition of patients with cystic fibrosis. J Cyst Fibros 12:497–503.

13. Maliniak ML, Stecenko AA, McCarty NA. 2016. A longitudinal analysis of chronic MRSA and *Pseudomonas aeruginosa* co-infection in cystic fibrosis: a single-center study. J Cyst Fibros 15:350–356.

14. Gangell C, Gard S, Douglas T, Park J, Klerk ND, Keil T, Brennan S, Ranganathan S, Robins-Browne R, Sly PD. 2011. Inflammatory responses to individual microorganisms in the lungs of children with cystic fibrosis. Clin Infect Dis 53:425–432.

15. Hendricks KJ, Burd TA, Anglen JO, Simpson AW, Christensen GD, Gainor BJ. 2001. Synergy between *Staphylococcus aureus* and *Pseudomonas aeruginosa* in a rat model of complex orthopaedic wounds. J Bone Joint Surg Am 83:855–861.

16. Dalton T, Dowd SE, Wolcott RD, Sun Y, Watters C, Griswold JA, Rumbaugh KP. 2011. An in vivo polymicrobial biofilm wound infection model to study interspecies interactions. PLoS One 6:e27317.

17. Pastar I, Nusbaum AG, Gil J, Patel SB, Chen J, Valdes J, Stojadinovic O, Plano LR, Tomic-Canic M, Davis SC. 2013. Interactions of methicillin resistant *Staphylococcus aureus* USA300 and *Pseudomonas aeruginosa* in polymicrobial wound infection. PLoS One 8:e56846.

18. Filkins LM, Graber JA, Olson DG, Dolben EL, Lynd LR, Bhuju S, O’Toole GA. 2015. Co-culture of *Staphylococcus aureus* with *Pseudomonas aeruginosa* drives *S. aureus* towards fermentative metabolism and reduced viability in a cystic fibrosis model. J Bacteriol 197:2252–2264.

19. Niggli S, Kümmerli R. 2020. Strain Background, species frequency, and environmental conditions are important in determining *Pseudomonas aeruginosa* and *Staphylococcus aureus* population dynamics and species coexistence. Appl Environ Microbiol 86:e00962–20.

20. Yung DBY, Sircombe KJ, Pletzer D. 2021. Friends or enemies? The complicated relationship between *Pseudomonas aeruginosa* and *Staphylococcus aureus*. Mol Microbiol 116:1–15.

21. Kessler E, Safrin M, Olson JC, Ohman DE. 1993. Secreted LasA of *Pseudomonas aeruginosa* is a staphylolytic protease. J Biol Chem 268:7503–7508.

22. Mashburn LM, Jett AM, Akins DR, Whiteley M. 2005. *Staphylococcus aureus* serves as an iron source for *Pseudomonas aeruginosa* during in vivo coculture. J Bacteriol 187:554–566.

23. Soberón-Chávez G, Lépine F, Déziel E. 2005. Production of rhamnolipids by *Pseudomonas aeruginosa*. Appl Microbiol Biotechnol 68:718–725.

24. Hoffman LR, Déziel E, D’Argenio DA, Lépine F, Emerson J, McNamara S, Gibson RL, Ramsey BW, Miller SI. 2006. Selection for *Staphylococcus aureus* small-colony variants due to growth in the presence of *Pseudomonas aeruginosa*. Proc Natl Acad Sci USA 103:19890–19895.

25. Harrison F, Paul J, Massey RC, Buckling A. 2008. Interspecific competition and siderophore-mediated cooperation in *Pseudomonas aeruginosa*. ISME J 2:49–55.

26. Lin J, Cheng J, Wang Y, Shen X. 2018. The *Pseudomonas* quinolone signal (PQS): Not just for quorum sensing anymore. Front Cell Infect Microbiol 8:230.

27. Nguyen AT, Oglesby-Sherrouse AG. 2016. Interactions between *Pseudomonas aeruginosa* and *Staphylococcus aureus* during co-cultivations and polymicrobial infections. Appl Microbiol Biotechnol 100:6141–6148.

28. Fazli M, Bjarnsholt T, Kirketerp-Møller K, Jørgensen B, Andersen AS, Krogfelt KA, Givskov M, Tolker-Nielsen T. 2009. Nonrandom distribution of *Pseudomonas aeruginosa* and *Staphylococcus aureus* in chronic wounds. J Clin Microbiol 47:4084–4089.

29. Phalak P, Chen J, Carlson RP, Henson MA. 2016. Metabolic modeling of a chronic wound biofilm consortium predicts spatial partitioning of bacterial species. BMC Syst Biol 10:90.

30. Tognon M, Köhler T, Gdaniec BG, Hao Y, Lam JS, Beaume M, Luscher A, Buckling A, van Delden C. 2017. Co-evolution with *Staphylococcus aureus* leads to lipopolysaccharide alterations in *Pseudomonas aeruginosa*. ISME J 11:2233–2243.

31. Dubern J-F, Diggle SP. 2008. Quorum sensing by 2-alkyl-4-quinolones in *Pseudomonas aeruginosa* and other bacterial species. Mol Biosyst 4:822–888.

32. Toyofuku M, Nakajima-Kambe T, Uchiyama H, Nomura N. 2010. The effect of a cell-to-cell communication molecule, *Pseudomonas* quinolone signal (PQS), produced by P. *aeruginosa* on other bacterial species. Microbes Environ 25:1–7.

33. Orazi G, O’Toole GA. 2017. *Pseudomonas aeruginosa* alters *Staphylococcus aureus* sensitivity to vancomycin in a biofilm model of cystic fibrosis infection. mBio 8:e00873–17.

34. Ramos AF, Woods DF, Shanahan R, Cano R, McGlacken GP, Serra C, O’Gara F, Reen FJ. 2020. A structure-function analysis of interspecies antagonism by the 2-heptyl-4-alkyl-quinolone signal molecule from *Pseudomonas aeruginosa*. Microbiology 166:169–179.

35. Bredenbruch F, Geffers R, Nimtz M, Buer J, Häussler S. 2006. The *Pseudomonas aeruginosa* quinolone signal (PQS) has an iron-chelating activity. Environ Microbiol 8:1318–1329.

36. Häussler S, Becker T. 2008. The pseudomonas quinolone signal (PQS) balances life and death in *Pseudomonas aeruginosa* populations. PLoS Biol 4:e1000166.

37. Abdalla MY, Hoke T, Seravalli J, Switzer BL, Bavitz M, Fliege JD, Murphy PJ, Britigan BE. 2017. Pseudomonas quinolone signal induces oxidative stress and inhibits heme oxygenase-1 expression in lung epithelial cells. Infect Immun 85:1–14.

38. Tognon M, Köhler T, Luscher A, Delden CV. 2019. Transcriptional profiling of *Pseudomonas aeruginosa* and *Staphylococcus aureus* during in vitro co-culture. BMC Genomics 20:30.

39. Clauditz A, Resch A, Wieland KP, Peschel A, Götz F. 2006. Staphyloxanthin plays a role in the fitness of *Staphylococcus aureus* and its ability to cope with oxidative stress. Infect Immun 74:4950–4953.

40. Antonic V, Stojadinovic A, Zhang B, Izadjoo MJ, Alavi M. 2013. *Pseudomonas aeruginosa* induces pigment production and enhances virulence in a white phenotypic variant of *Staphylococcus aureus*. Infect Drug Resist 6:175–186.

41. Biswas L, Biswas R, Schlag M, Bertram R, Götz F. 2009. Small-colony variant selection as a survival strategy for *Staphylococcus aureus* in the presence of *Pseudomonas aeruginosa*. Appl Environ Microbiol 75:6910–6912.

42. Tuchscherr L, Medina E, Hussain M, Völker W, Heitmann V, Niemann S, Holzinger D, Roth J, Proctor RA, Becker K, Peters G, Löffler B. 2011. *Staphylococcus aureus* phenotype switching: An effective bacterial strategy to escape host immune response and establish a chronic infection. EMBO Mol Med 3:129–141.

43. Kahl BC, Becker K, Löffler B. 2016. Clinical significance and pathogenesis of staphylococcal small colony variants in persistent infections. Clin Microbiol Rev 29:401–427.

44. Burnside K, Lembo A, de Los Reyes M, Iliuk A, Binhtran NT, Connelly JE, Lin WJ, Schmidt BZ, Richardson AR, Fang FC, Tao WA, Rajagopal L. 2010. Regulation of hemolysin expression and virulence of *Staphylococcus aureus* by a serine/threonine kinase and phosphatase. PLoS One 5:e11071.

45. Shopsin B, Drlica-Wagner A, Mathema B, Adhikari RP, Kreiswirth BN, Novick RP. 2008. Prevalence of agr dysfunction among colonizing *Staphylococcus aureus* strains. J Infect Dis 198:1171–1174.

46. Traber KE, Lee E, Benon S, Corrigan R, Cantera M, Shopsin B, Novick RP. 2008. agr function in clinical *Staphylococcus aureus* isolates. Microbiology 154:2265–2274.

47. Zeden MS, Kviatkovski I, Schuster CF, Thomas VC, Fey PD, Gründling A. 2020. Identification of the main glutamine and glutamate transporters in *Staphylococcus aureus* and their impact on c-di-AMP production. Mol Microbiol 113:1085–1100.

48. Ibberson CB, Whiteley M. 2020. The social life of microbes in chronic infection. Curr Opin Microbiol 53:44–50.

49. Rezzoagli C, Granato ET, Kümmerli R. 2020. Harnessing bacterial interactions to manage infections: a review on the opportunistic pathogen *Pseudomonas aeruginosa* as a case example. J Med Microbiol 69:147–161.

50. Serra R, Grande R, Butrico L, Rossi A, Settimio UF, Caroleo B, Amato B, Gallelli L, Franciscis SD. 2015. Chronic wound infections: The role of *Pseudomonas aeruginosa* and *Staphylococcus aureus*. Expert Rev Anti Infect Ther 13:605–613.

51. Camus L, Briaud P, Vandenesch F, Moreau K. 2021. How bacterial adaptation to cystic fibrosis environment shapes interactions between *Pseudomonas aeruginosa* and *Staphylococcus aureus*. Front Microbiol 12:617784.

52. Diggle SP, Matthijs S, Wright VJ, Fletcher MP, Chhabra SR, Lamont IL, Kong X, Hider RC, Cornelis P, Camara M, Williams P. 2007. The *Pseudomonas aeruginosa* 4-quinolone signal molecules HHQ and PQS play multifunctional roles in quorum sensing and iron entrapment. Chem Biol 14:87–96.

53. Popat R, Harrison F, da Silva AC, Easton SA, McNally L, Williams P, Diggle SP. 2017. Environmental modification via a quorum sensing molecule influences the social landscape of siderophore production. Proc R Soc Lond B 284:20170200.

54. Diep BA, Gill SR, Chang RF, Phan TH, Chen JH, Davidson MG, Lin F, Lin J, Carleton HA, Mongodin EF, Sensabaugh GF, Perdreau-Remington F. 2006. Complete genome sequence of USA300, an epidemic clone of community-acquired meticillin-resistant *Staphylococcus aureus*. Lancet 367:731–739.

55. Pollitt EJG, West SA, Crusz SA, Burton-Chellew MN, Diggle SP. 2014. Cooperation, quorum sensing, and evolution of virulence in *Staphylococcus aureus*. Infect Immun 82:1045–1051.

56. Cheung GYC, Bae JS, Otto M. 2021. Pathogenicity and virulence of *Staphylococcus aureus*. Virulence 12:547–569.

57. Huemer M, Mairpady Shambat S, Brugger SD, Zinkernagel AS. 2020. Antibiotic resistance and persistence-Implications for human health and treatment perspectives. EMBO Rep 21:e51034.

58. Schieber M, Chandel NS. 2014. ROS function in redox signaling and oxidative stress. Curr Biol 24:R453–R462.

59. Deurenberg RH, Stobberingh EE. 2008. The evolution of *Staphylococcus aureus*. Infect Genet Evol 8:747–763.

60. Corrigan RM, Abbott JC, Burhenne H, Kaever V, Gründling A. 2011. c-di-AMP is a new second messenger in *Staphylococcus aureus* with a role in controlling cell size and envelope stress. PLoS Pathog 7:e1002217.

61. Lensmire JM, Dodson JP, Hsueh BY, Wischer MR, Delekta PC, Shook JC, Ottosen EN, Kies PJ, Ravi J, Hammer ND. 2020. The *Staphylococcus aureus* cystine transporters TcyABC and TcyP facilitate nutrient sulfur acquisition during Infection. Infect Immun 88:e00690–19.

62. Zeden MS, Burke O, Vallely M, Fingleton C, O’Gara JP. 2021. Exploring amino acid and peptide transporters as therapeutic targets to attenuate virulence and antibiotic resistance in *Staphylococcus aureus*. PLoS Pathog 17:e1009093.

63. Niggli S, Wechsler T, Kümmerli R. 2021. Single-cell imaging reveals that *Staphylococcus aureus* Is highly competitive against *Pseudomonas aeruginosa* on surfaces. Front Cell Infect Microbiol 11:733991.

64. Hall JW, Yang J, Guo H, Ji Y. 2017. The *Staphylococcus aureus* AirSR two-component system mediates reactive oxygen species resistance via transcriptional regulation of staphyloxanthin production. Infect Immun 85:1–12.

65. Bolger AM, Lohse M, Usadel B. 2014. Trimmomatic: a flexible trimmer for Illumina sequence data. Bioinformatics 30:2114–2120.

