## Supplemental Material for "Rapid and strain-specific resistance evolution of *Staphylococcus aureus* against inhibitory molecules secreted by *Pseudomonas aeruginosa*"

**Supplemental Figure S1**

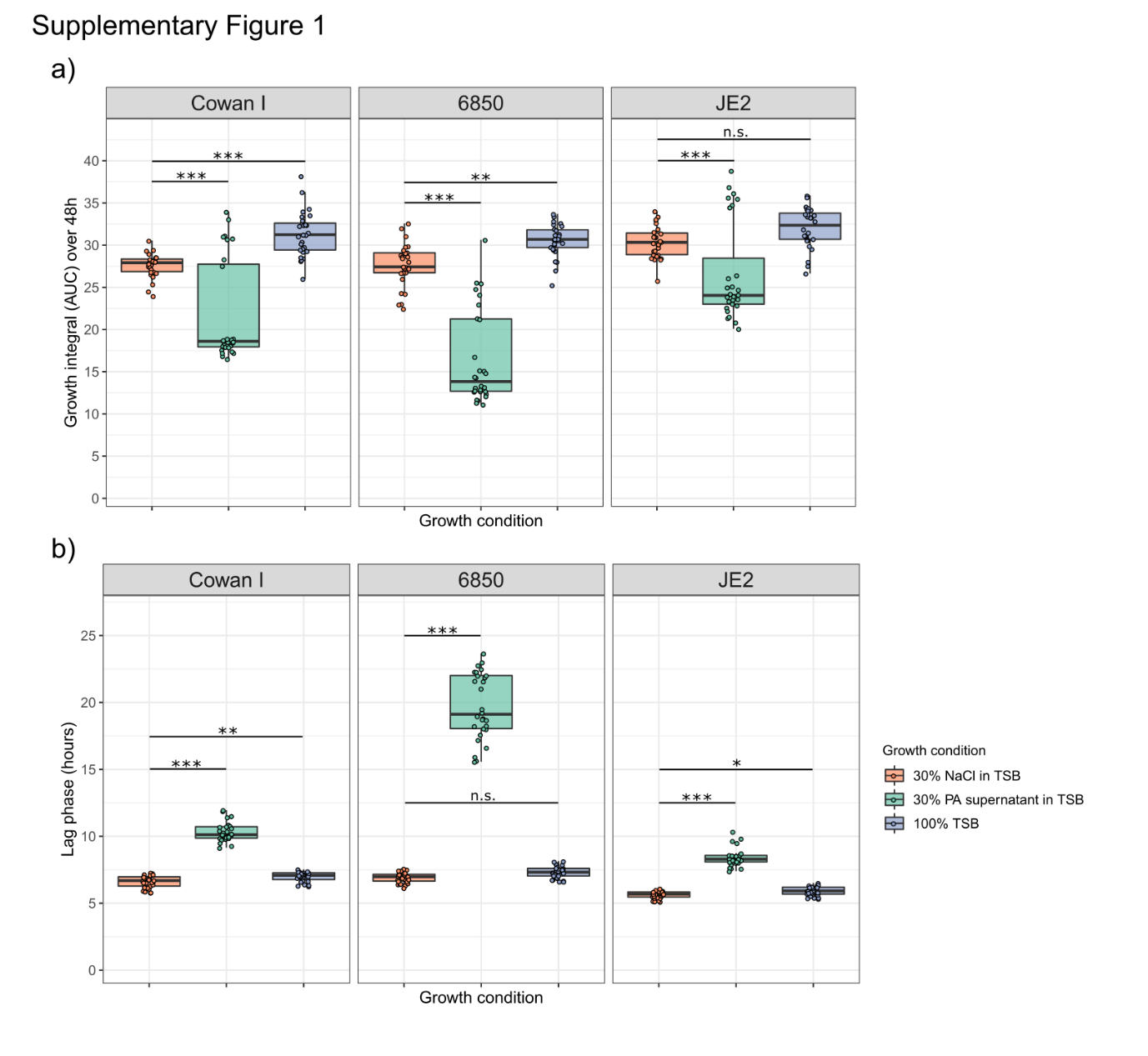
**Figure S1.** Quantification of growth (growth integral and lag phase) of the three SA strains Cowan I, 6850 and JE2 under the three different supernatant assay conditions. a) Growth integrals (area under the curve, AUC) over 48 hours at 37 °C under the three different growth conditions. b) Lag phase (in hours) under the three different growth conditions. There is some variability in the growth integrals in 30% PA supernatant + TSB, and particularly in two out of seven experiments, all three SA strains reached, although initially inhibited, similar growth integrals as the controls. In contrast, there was little variation in the shifts in lag phase, which were always consistently extended in the presence of PA supernatant across all experiments and for all three SA strains. The box plots show the median (bold line) with the first and third quartiles. The whiskers cover the 1.5* inter-quartile range (IQR) or extend from the lowest to the highest value if they fall within the 1.5* IQR. Data is from seven independent experiments with 28 replicates per SA strain per growth condition in total. Asterisks denote significant differences from the control (30% NaCl in TSB) as determined by Tukey’s HSD with adjusted p values. *** p < 0.001; ** p < 0.01; * p < 0.05; n.s. not significant. The results from all statistical analyses can be found in the Statistics Source File.

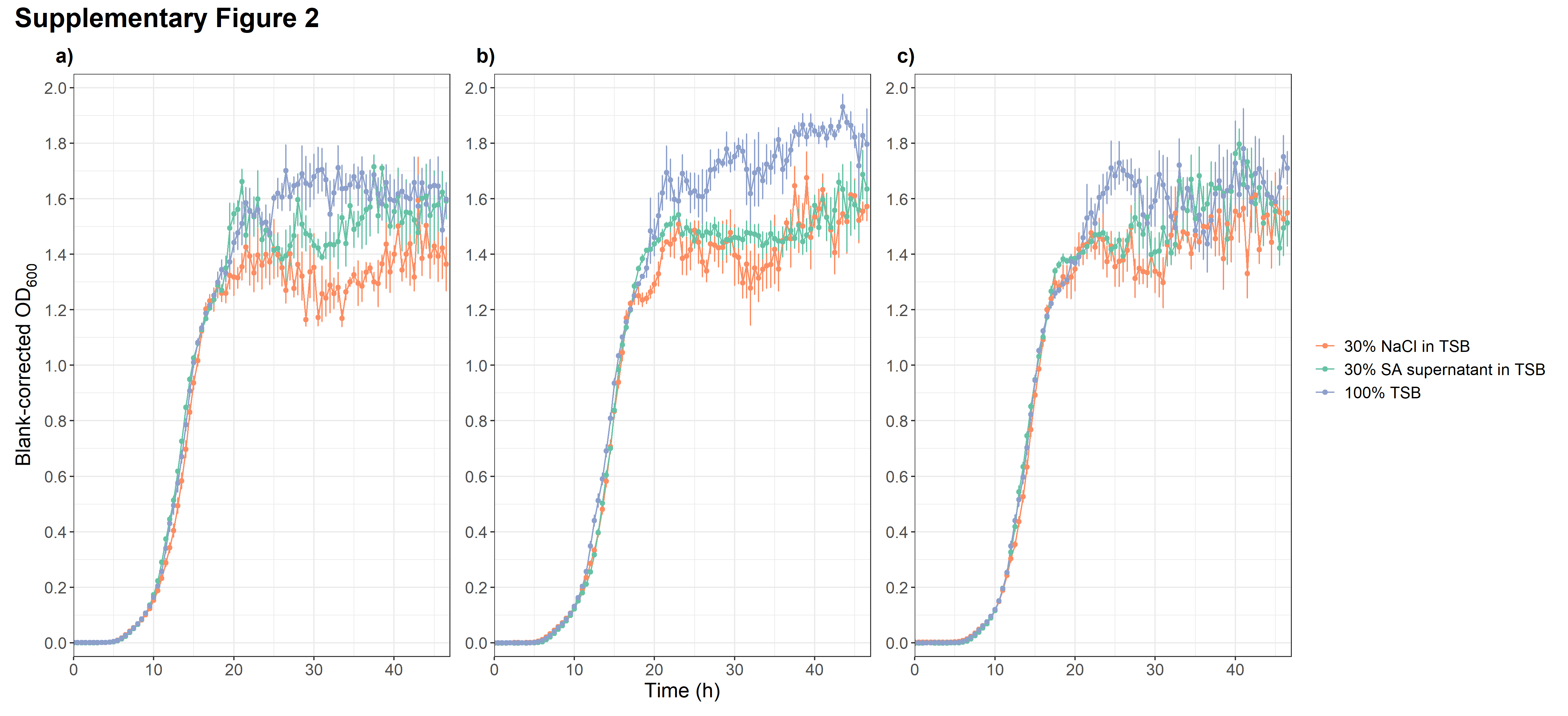
 **Supplemental Figure S2**

**Figure S2.** Supernatants of SA strains do not inhibit growth of PA. Growth of PA strain PAO1 was measured in the presence of a) Cowan I supernatant; b) 6850 supernatant; and c) JE2 supernatant, over 48 hours at 37°C. As controls, we used 30% NaCl in TSB and 100% TSB. None of the SA supernatants were able to inhibit growth of PA. Data is from one experiment per SA supernatant with four replicates per condition. Error bars show standard errors.

**Supplemental Figure S3**

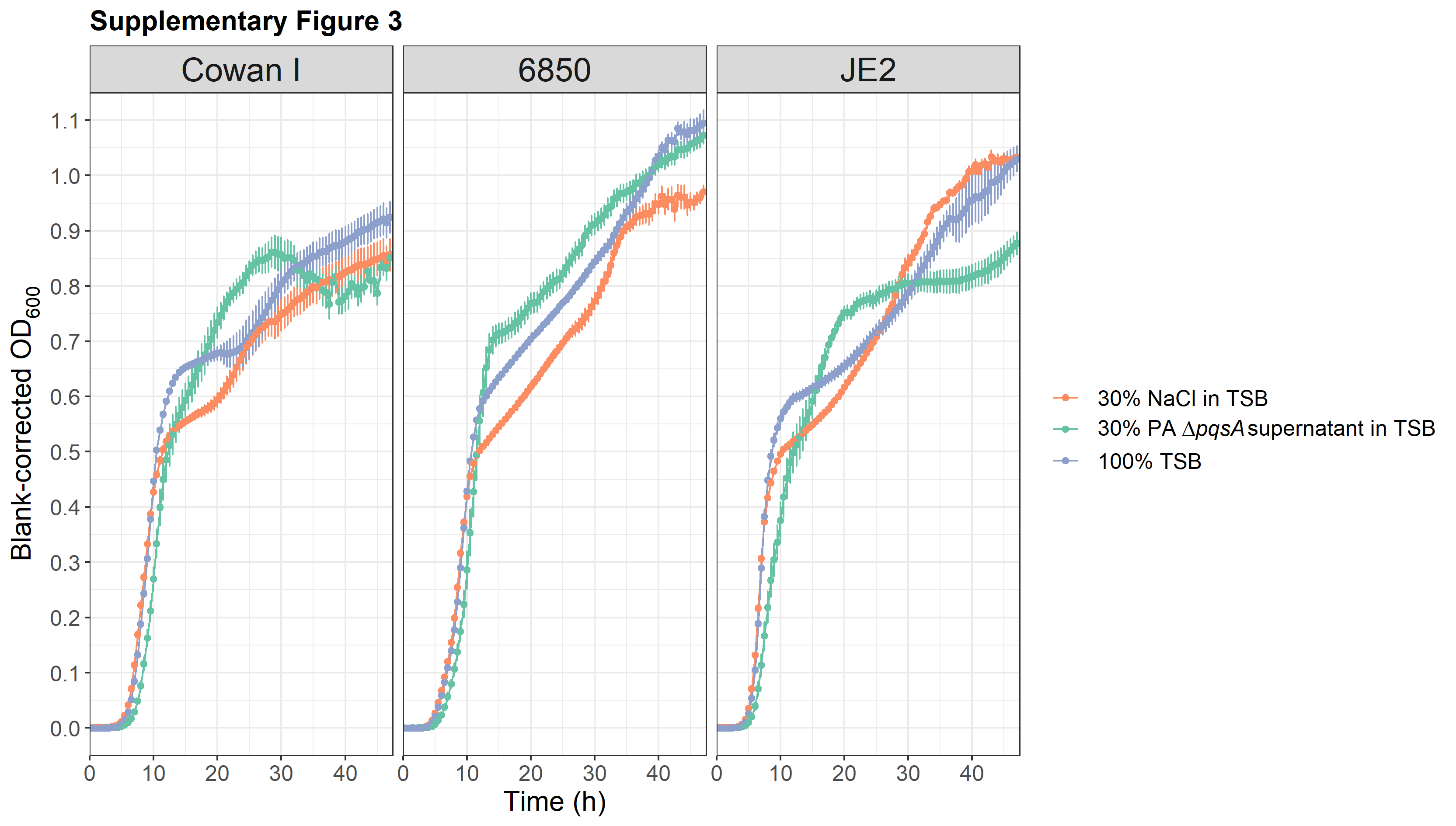
**Figure S3.** Supernatant of PAΔ*pqsA* does not inhibit growth of SA anymore. Growth of SA strains Cowan I, 6850 and JE2 was recorded by measuring OD_600_ over 48 hours at 37 °C. As controls, we used growth in 30% NaCl in TSB and 100% TSB. Data is from two independent experiments with eight replicates per SA strain per condition in total. Error bars show standard error.

**Supplemental Figure S4**

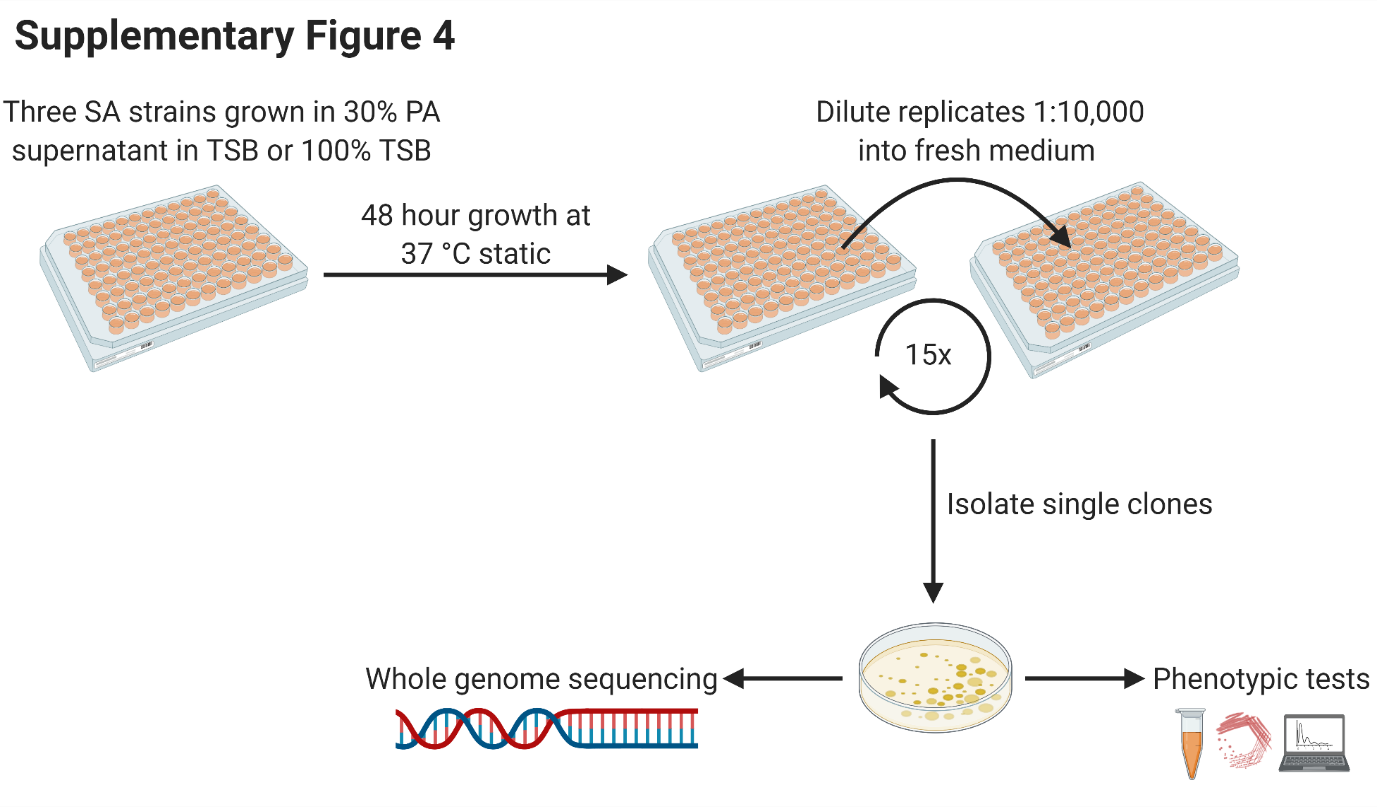

**Figure S4.** Workflow for the experimental evolution of SA strains Cowan I, 6850 and JE2 in the presence or absence of PA supernatant. The experimental evolution consisted of two treatments, 100% TSB (control) and 30% PA supernatant + 70% TSB (condition of interest). We evolved seven independent replicates per SA strain per treatment for one month at 37 °C in a static incubator by transferring growing cultures to fresh medium every 48 hours. Prior to transfers, OD_600_ values of all cultures were recorded. After transfers, cultures in the old plates were frozen as glycerol stocks at -80 °C. At the end of the evolution experiment, we isolated five clones per replicate, resulting in 210 individual clones from 42 different populations. 150 clones from 30 populations were then used for in-depth phenotypic and genetic characterization. All clones were maintained individually as glycerol stocks at -80 °C.

**Supplemental Figure S5**

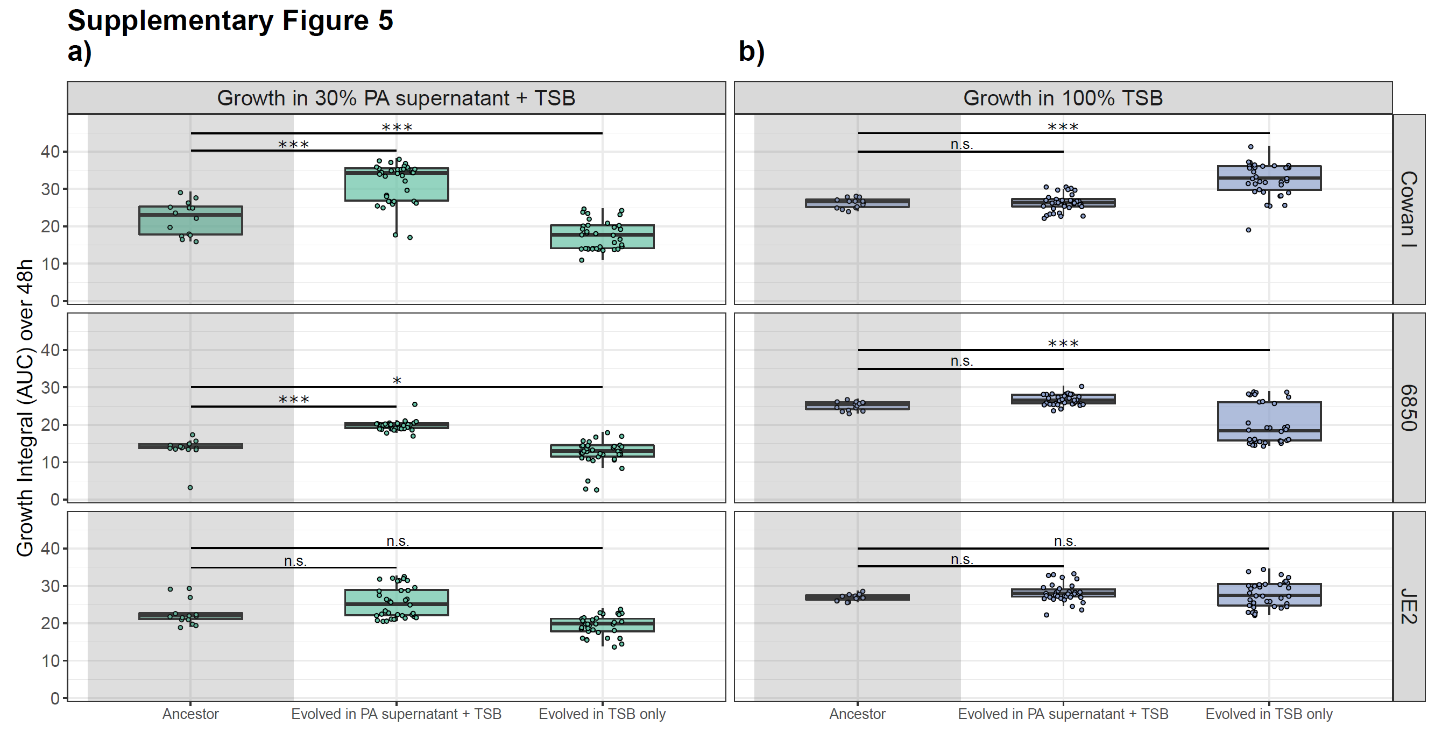

**Figure S5.** Most clones from Cowan I, 6850 and JE2 evolved resistance to inhibitory molecules produced by PA when exposed to its supernatant during experimental evolution (depicted as growth integrals, area under the curve). Clones were isolated from populations of SA strains either evolved in 30% PA supernatant + 70% TSB or 100% TSB over 30 days, with transfer to fresh medium every 48 hours. a) Growth of ancestral strain (grey-shaded area) and evolved clones in medium containing 30% PA supernatant + 70% TSB. SA clones from Cowan I and 6850 evolved in the presence of PA supernatant significantly improved growth, while all clones evolved in TSB alone are still inhibited by the PA supernatant. b) Growth of ancestral strain (grey-shaded area) and evolved clones in 100% TSB medium. SA clones evolved in the presence of PA supernatant did not improve growth in TSB alone, while many clones evolved in TSB did so for two of the three SA strains (except 6850). Box plots show the median (bold line), the first and third quartiles, and the 1.5* inter-quartile range (IQR, whiskers) or the range from the lowest to highest value if all values fall within the 1.5* IQR. Asterisks denote significant differences between the ancestral strain and evolved clones. *** p < 0.001; * p < 0.05; n.s. not significant. P-values are adjusted by the false discovery rate method.

**Supplemental Figure S6**

a) **b)**

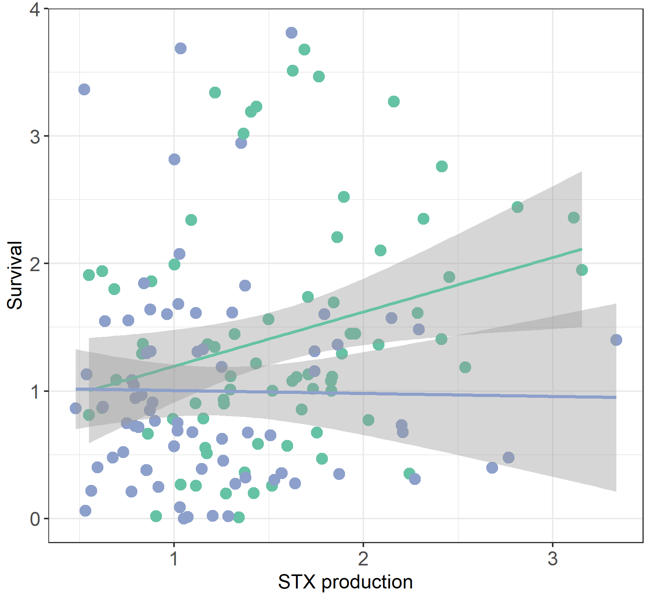

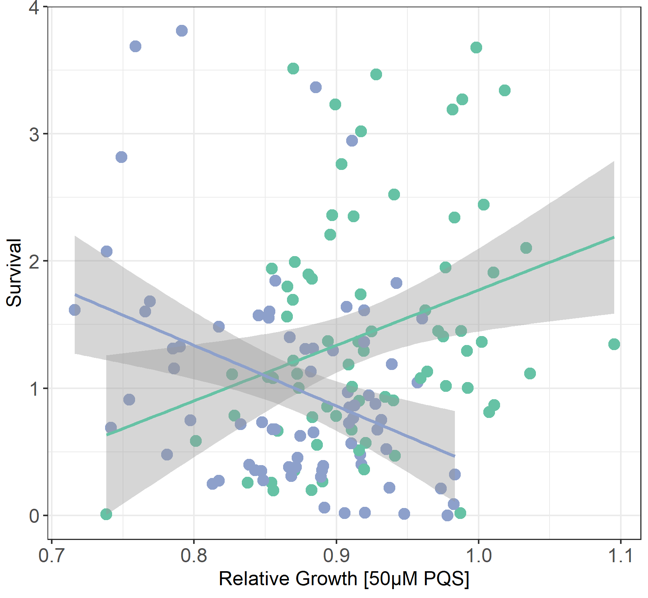

**Figure S6.** Relationship between survival in the presence of hydrogen peroxide H_2_O_2_ and (a) growth under PQS exposure and (b) staphyloxanthin (STX) production for clones evolved in 30% PA supernatant + 70% TSB (green dots) and clones evolves in TSB alone (blue dots). For clones evolved in PA supernatant, there were positive correlations between the two variables (H_2_O_2_ survival versus PQS growth, Pearson’s product-moment correlation: r_73_ = 0.30, p = 0.0092; H_2_O_2_ survival versus STX production: r_73_ = 0.27, p = 0.0206). For clones evolved in TSB alone, there was a negative correlation between H_2_O_2_ survival and PQS growth (r_73_ = -0.37, p = 0.0010) and no significant correlation between H_2_O_2_ survival and STX production (r_73_ = -0.02, p = 0.8920).

**Supplemental Figure S7**

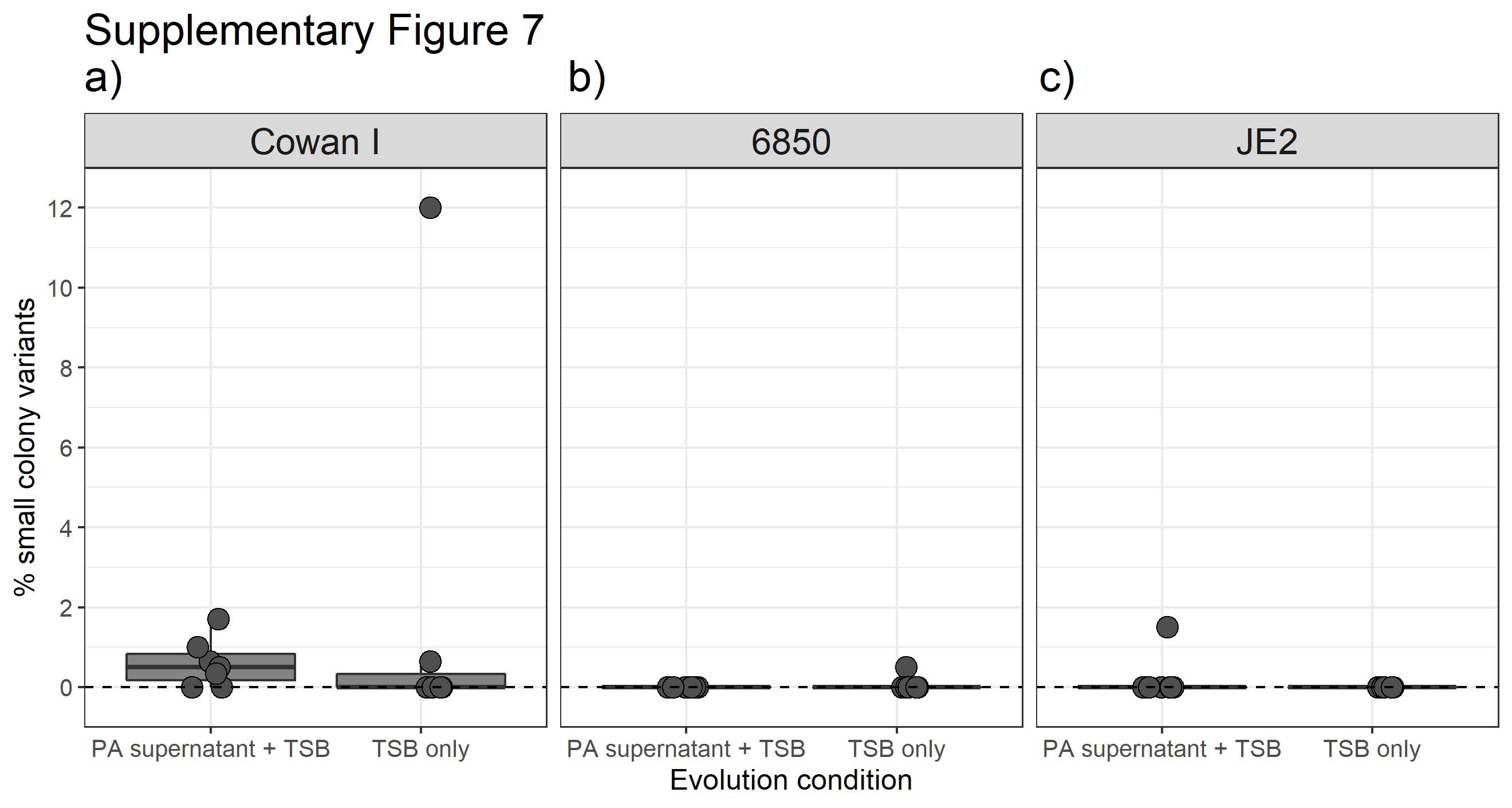

**Figure S7.** Five out of seven populations from Cowan I evolved in PA supernatant + TSB have an elevated percentage of small colony variants (SCVs). Shown are the percentages of SCVs by SA population evolved in either PA supernatant + TSB or TSB alone for the three SA strains a) Cowan I, b) 6850 and c) JE2. The datapoints shown are the average from two independent counting assays. As populations were sub-cultured twice in TSB before counting SCVs, most unstable SCVs may have reverted until that stage, and we likely only counted stable SCVs in this assay. In this plot, all the populations are shown (seven populations per SA strain and evolution condition). The box plots show the median (bold line) with the first and third quartiles. The whiskers cover the 1.5* inter-quartile range (IQR) or extend from the lowest to the highest value if they fall within the 1.5* IQR. The dotted line at zero represents the ancestral state.

**Supplemental Figure S8**

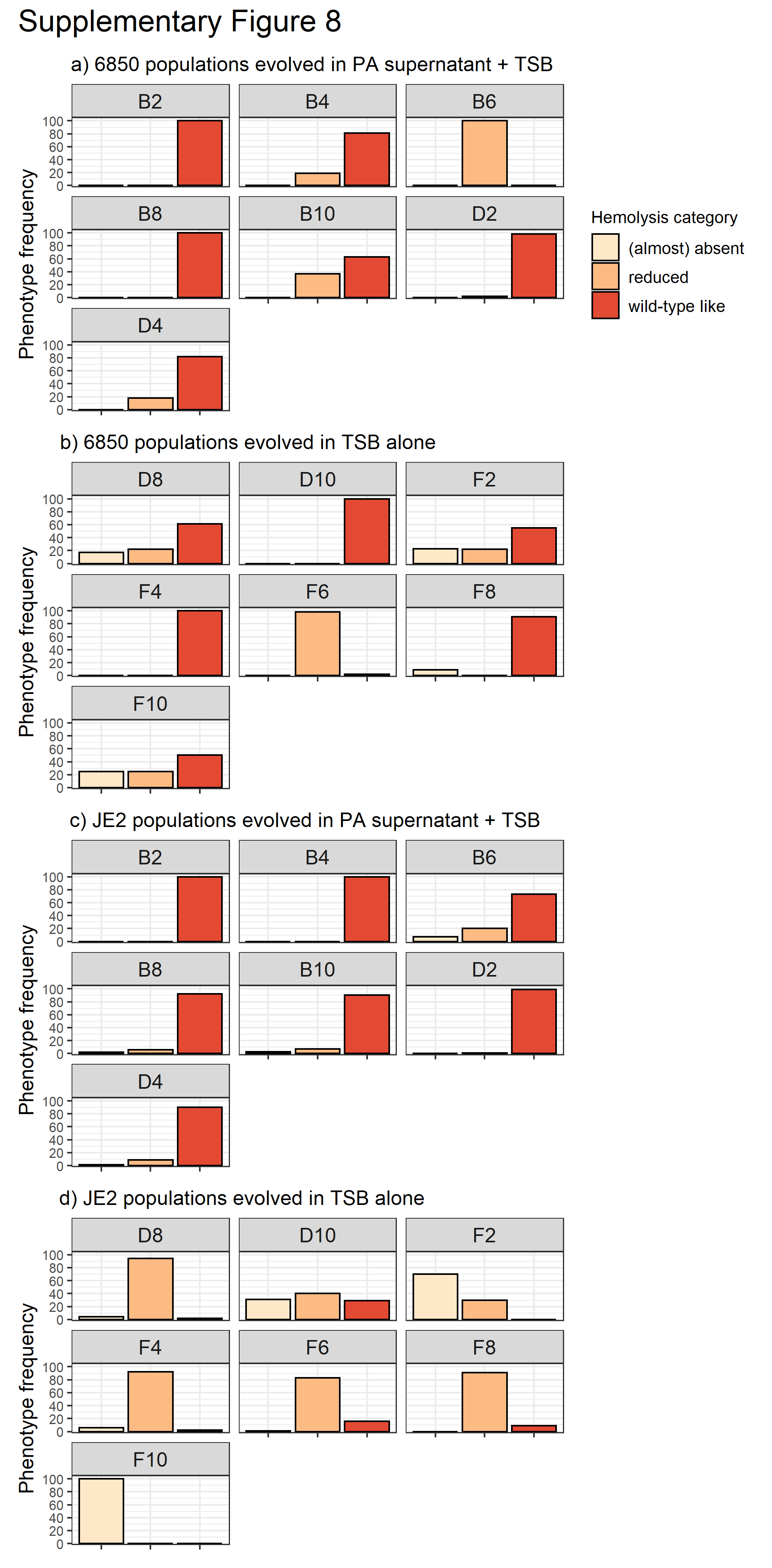

**Figure S8.** Frequency of three hemolysis phenotypes in populations of a) 6850 evolved in PA supernatant + TSB; b) 6850 evolved in TSB alone; c) JE2 evolved in PA supernatant + TSB; d) JE2 evolved in TSB alone. Evolved populations were diluted, plated on sheep blood agar plates and hemolysis patterns of growing colonies were classified into three categories: (almost) absent, reduced, wild-type like. For 6850, the frequency of the three hemolysis phenotypes was similar across populations and evolution conditions a) and b), with most of the colonies expressing wild-type like phenotypes. This is not the case for JE2 populations. Here, the frequency of the three hemolysis phenotypes is markedly different between the evolution conditions c) and d). Specifically, hemolysis levels are greatly reduced in populations evolved in TSB alone, while wild-type like phenotypes are maintained in the presence of PA supernatant. These data are consistent with the data for the individual clones analyzed in Figure 6.

**Supplemental Table S1.** Strains used for this study.

| **Species and strain name** | **Origin** | **Description** | **Reference** |
| --- | --- | --- | --- |
| ***Pseudomonas aeruginosa* (PA)** |  |  |  |
| PAO1 | Wound | Commonly used *Pseudomonas aeruginosa* laboratory strain. | ATCC 15692 |
| ***Staphylococcus aureus* (SA)** |  |  |  |
| Cowan I | Septic arthritis | MSSA isolate. Highly invasive, but not cytotoxic. Agr-defective. | ATCC 12598 |
| 6850 | Osteomyelitis | MSSA isolate. Highly invasive, cytotoxic, and hemolytic. | ATCC 53657 |
| JE2 | Skin and soft tissue infection | USA300 CA-MRSA isolate. Highly virulent, cytotoxic, and hemolytic. | NARSA |

CA-MRSA: Community-acquired methicillin-resistant *S. aureus*

MSSA: Methicillin-sensitive *S. aureus*

Agr: Accessory gene regulator

**Supplemental Table S2**

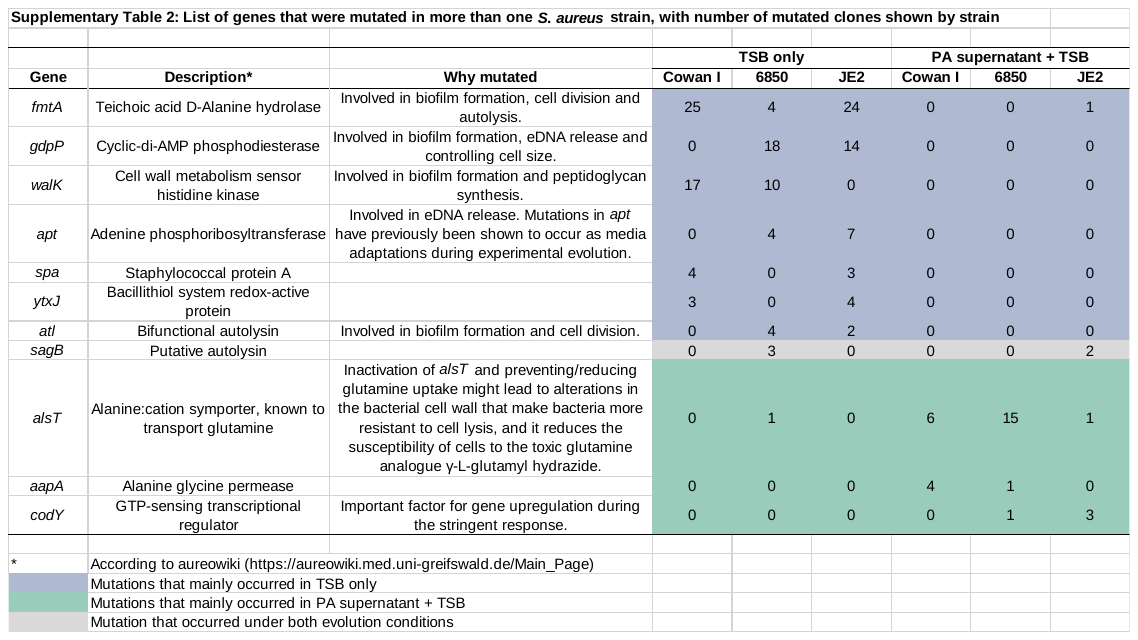
